# CUPID-seq enables highly multiplexed amplicon sequencing via combinatorial in-line dual indexing

**DOI:** 10.64898/2026.05.20.726713

**Authors:** Beverly Fu, Rachel L. Porter, Handuo Shi, Allison C. Ea, Ana M. Espeleta, Achuthan Ambat, David A. Relman, Kerwyn Casey Huang, Katherine S. Xue

## Abstract

Targeted amplicon sequencing is widely used to profile genetic variation in defined genomic regions. In microbial ecology, for example, amplicon sequencing of the 16S and 18S ribosomal RNA genes has been transformative for characterizing microbial communities. However, on high-capacity sequencing platforms with patterned flow cells, throughput is constrained by the requirement for unique dual indexes (UDIs), which increases primer costs and limits the number of samples that can be pooled per sequencing run. Here, we introduce CUPID-seq (<u>C</u>ombinatorial, <u>U</u>nique, <u>P</u>hased, In-line <u>D</u>ual-indexed <u>seq</u>uencing), a highly multiplexed amplicon-sequencing strategy that increases scalability through combinatorial indexing across two rounds of PCR. CUPID-seq introduces phased, in-line UDIs during Round 1 gene-specific amplification, enabling multiple samples to share the same Illumina UDI during Round 2 PCR while remaining uniquely identifiable. This design reduces upfront costs by up to 85% and reduces library preparation time and reagent use by up to 40%. We develop and validate CUPID-seq primers targeting the 16S V4 region and provide a computational workflow for demultiplexing in-line indexes. Although optimized here for 16S-based profiling, CUPID-seq can be readily adapted to other user-defined amplicons. By reducing cost and increasing multiplexing capacity, CUPID-seq enables users to leverage high-throughput sequencing platforms more effectively across diverse biological contexts.

## Introduction

Targeted amplicon sequencing has transformed the study of genetic variation by enabling rapid, cost-effective characterization of genetic diversity at defined genomic regions of interest. By amplifying short (<1 kB), locus-specific sequences, researchers have developed scalable approaches to perform functional screens^1–3^, track evolving lineages^4^, and detect patient-specific tumor mutations^5^, among many other applications. In microbial ecology, amplicon sequencing of ribosomal RNA (rRNA) genes, including the 16S rRNA gene^6–10^, 18S rRNA gene^11,12^, and internal transcribed spacer (ITS)^13,14^ regions, has enabled cultivation-independent profiling of community composition. These methods form the basis of DNA metabarcoding approaches that identify organisms directly from environmental or bulk samples by amplifying universal barcode loci^15^. Because of these widespread applications, improvements that increase throughput, reduce cost, and streamline analysis can accelerate progress across diverse areas of genomics, evolutionary biology, and microbial ecology.

Recent advances in Illumina sequencing technology, particularly the introduction of patterned flow cells, have dramatically increased sequencing capacity while reducing per-base cost. Modern platforms can generate hundreds of millions of reads per run, theoretically enabling thousands of amplicon libraries to be sequenced simultaneously. However, microbiome laboratories are often unable to fully exploit this expanded capacity. On patterned flow cell platforms, samples must be labeled with unique dual indexes (UDIs) to mitigate index hopping^16^, which constrains the number of libraries that can be multiplexed and substantially increases primer costs. As a result, the practical throughput of amplicon sequencing is frequently limited not by sequencing output, but by indexing architecture.

Here, we introduce CUPID-seq (<u>C</u>ombinatorial, <u>U</u>nique, <u>P</u>hased, In-line <u>D</u>ual-indexed <u>seq</u>uencing), a highly multiplexed amplicon-sequencing strategy designed to overcome this bottleneck. CUPID-seq modifies standard two-step PCR workflows by introducing phased, in-line UDIs during the first round of gene-specific amplification (Round 1; UDI-1). These Round 1 indexes are combined with standard Illumina UDIs during a second PCR (Round 2; UDI-2), enabling combinatorial scaling while maintaining compatibility with patterned flow cells. This design allows multiple samples to share the same Round 2 UDI while remaining uniquely identifiable, reducing primer costs by up to 85% compared to conventional approaches and enabling the simultaneous sequencing of thousands of amplicon libraries on a single Illumina run.

We develop and validate CUPID-seq primers targeting the V4 region of the 16S rRNA gene and demonstrate that sequencing results are largely consistent with established Earth Microbiome Project (EMP) primers^8,9^. We further show that Round 1 PCR products can be pooled prior to Round 2 PCR under appropriate conditions, reducing library preparation time and reagent use by up to 40%. A complementary computational workflow enables demultiplexing of phased in-line indexes and supports straightforward implementation. Although optimized here for 16S rRNA gene-based microbiome profiling (16S sequencing), the CUPID-seq framework can be readily adapted to other genomic loci. We anticipate that this protocol will provide a flexible and scalable approach for high-throughput amplicon sequencing that broadens access to modern sequencing capacity across biological disciplines.

### Limitations of current approaches

The introduction of patterned flow cells has substantially increased the throughput and reduced the per-base cost of Illumina sequencing (**Table 1**)^17,18^. Earlier instruments such as the MiSeq v3 platform use non-patterned flow cells and produce ∼20 million reads per run at a cost of approximately $100 per million reads. More recently introduced benchtop instruments have transitioned to patterned flow-cell technology and can generate 5–25 million reads per run (MiSeq i100) or 5–100 million reads per run (MiSeq i100 Plus), depending on the selected reagent kit. At larger scales, newer platforms including the NextSeq 1000/2000 and NovaSeq 6000 also use patterned flow cells and generate 100–800 million reads per lane at ∼$7–15 per million reads. Together, these advances represent an order-of-magnitude reduction in per-base cost and a dramatic expansion of read capacity.

**Table 1:**
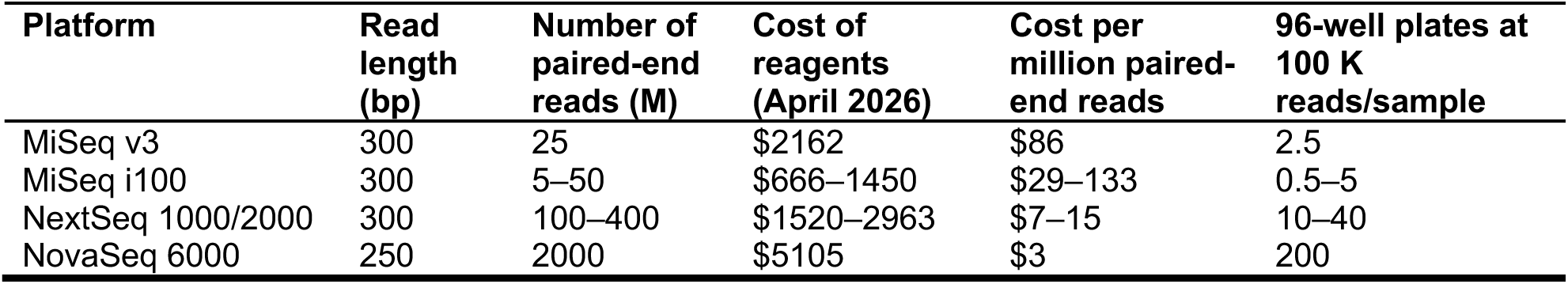
Cost and throughput of current Illumina sequencing platforms.

| Platform | Read length (bp) | Number of paired-end reads (M) | Cost of reagents (April 2026) | Cost per million paired-end reads | 96-well plates at 100 K reads/sample |
| --- | --- | --- | --- | --- | --- |
| MiSeq v3 | 300 | 25 | \$2162 | \$86 | 2.5 |
| MiSeq i100 | 300 | 5–50 | \$666–1450 | \$29–133 | 0.5–5 |
| NextSeq 1000/2000 | 300 | 100–400 | \$1520–2963 | \$7–15 | 10–40 |
| NovaSeq 6000 | 250 | 2000 | \$5105 | \$3 | 200 |

For amplicon sequencing applications, however, this expanded capacity is not fully accessible. On patterned flow cell platforms, libraries must use unique dual indexes (UDIs) to mitigate index hopping^16,17,19,20^, in which reads are assigned to the wrong forward or reverse index. Because each sample requires a distinct pair of UDIs, the number of libraries that can be pooled on a single run is limited by the number of available index pairs. As of 2026, only 384 UDIs are currently available from Illumina^21^, establishing an upper bound on multiplexing without custom primer design. Designing and synthesizing additional UDIs typically costs $10–15 per index, making large-scale multiplexing prohibitively expensive in practice. Consequently, while modern sequencers are capable of generating hundreds of millions of reads (sufficient for thousands of 16S libraries with sequencing depths of ∼100,000 reads per sample), the practical throughput of pooled amplicon sequencing is constrained by indexing architecture and primer cost rather than sequencing output.

These limitations create a mismatch between sequencing capacity and indexing scalability, particularly for microbiome studies that routinely process hundreds to thousands of samples. A strategy that increases the number of distinguishable libraries without requiring a proportional expansion in UDI primer sets would enable laboratories to more fully exploit modern high-throughput sequencing platforms.

### Overview of protocol

To increase the throughput of amplicon sequencing, we modified a standard two-step PCR workflow for generating indexed amplicon libraries^22^ (**Fig. 1**). In conventional protocols, the first PCR (Round 1) amplifies the genomic region of interest using primers that include a gene-specific sequence and an overhang compatible with sequencing adapters. The second PCR (Round 2) attaches flow-cell adapters and a UDI pair that distinguishes each sample within a sequencing run^22^. Because each sample must be labeled with a distinct pair of forward and reverse UDIs located on the Round 2 primers^20,21^, one unique UDI primer pair is required per sample (**Fig. 1A**).

**Figure 1:**
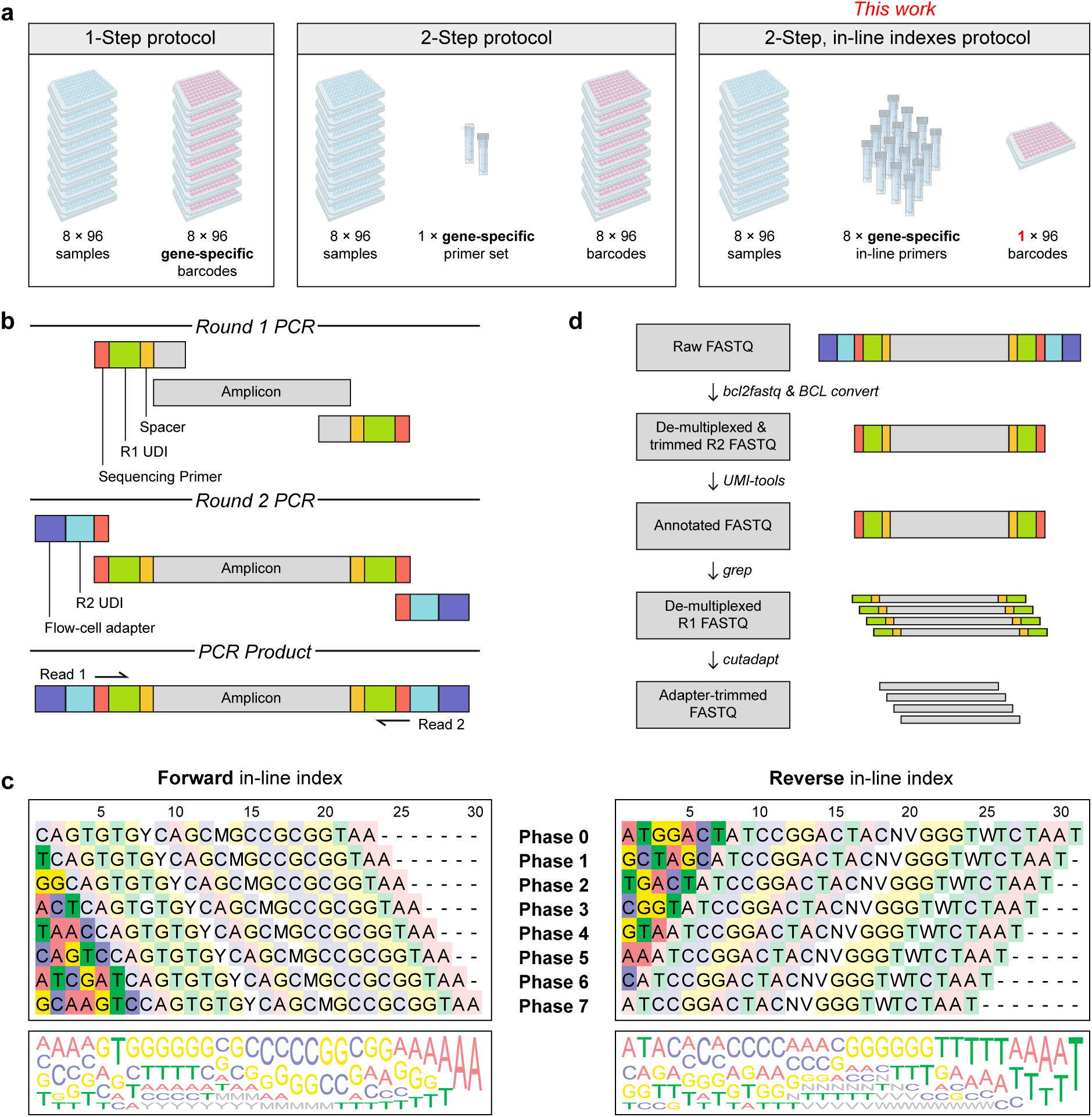
CUPID-seq uses phased, in-line unique dual indexes (UDIs) to increase the throughput and decrease the cost of amplicon sequencing. A) Comparison of the primers needed to prepare 8×96 = 768 samples using 1-step, 2-step, and 2-step with in-line indexes (CUPID-seq) protocols. The 1-step protocol requires 768 sets of unique single indexes for each gene of interest. The 2-step protocol uses a Round 1 gene-specific primer set to amplify the gene of interest and then 768 Round 2 UDIs. The CUPID-seq protocol uses 8 unique Round 1 gene-specific primer sets to create base diversity and decrease the number of required Round 2 UDIs to 96. B) Schematic of the components incorporated during CUPID-seq library generation. In the Round 1 PCR, the region of interest is amplified using primers that contain a spacer region (yellow), phased in-line UDI (green), and Illumina Nextera sequencing primer overhang (red). In the Round 2 PCR, the Round 1 PCR product is further amplified with primers that contain UDI barcodes (teal) and Illumina flow-cell adapters (blue). C) The 8 pairs of phased primers create sufficient base diversity so that pooled amplicon libraries can be sequenced using high-throughput Illumina two-color chemistries with minimum PhiX spike-in. Deeply hued boxes represent the forward and reverse in-line indexes. Bottom: the sequence logo shows the nucleotide diversity at each position. D) Computational workflow to demultiplex reads into individual samples.

CUPID-seq modifies this architecture by introducing phased, in-line UDIs during Round 1 amplification (**Fig. 1B**). We developed eight sets of Round 1 UDIs consisting of seven total nucleotides distributed between the forward and reverse primers (Phase-0 to Phase-7). For example, the Phase-2 primer pair contains a 2-bp forward and 5-bp reverse index, whereas the Phase-4 primer pair contains a 4-bp forward and 3-bp reverse index (**Fig. 1C**). The first 7 bp of each forward and reverse primer, which include the Round 1 UDI and the downstream gene-specific region, differ from all other primers by at least three nucleotides to reduce misassignment due to sequencing errors. These in-line indexes are positioned between the sequencing primer and the gene-specific region, ensuring that they are sequenced at the start of each read.

To minimize amplification bias between Round 1 primer sets, a constant 4-bp spacer sequence (forward: CAGT; reverse: ATCC) was inserted between the gene-specific region and the in-line index. This spacer incorporates the 2-bp pad sequences (forward: GT; reverse: CC) from the EMP primers^9^ adjacent to the gene-specific primer regions, maintaining interoperability with common 16S sequencing workflows.

The phased, in-line architecture addresses two limitations of conventional indexing strategies (**Table 2**). First, the phased distribution of index lengths introduces nucleotide diversity at the beginning of the sequencing reads, reducing the need for additional PhiX DNA or shotgun libraries^23,24^ (**Fig. 1C**). Second, and more importantly, Round 1 UDIs can be combined with any Round 2 UDI, enabling combinatorial scaling of distinguishable libraries.

**Table 2:**
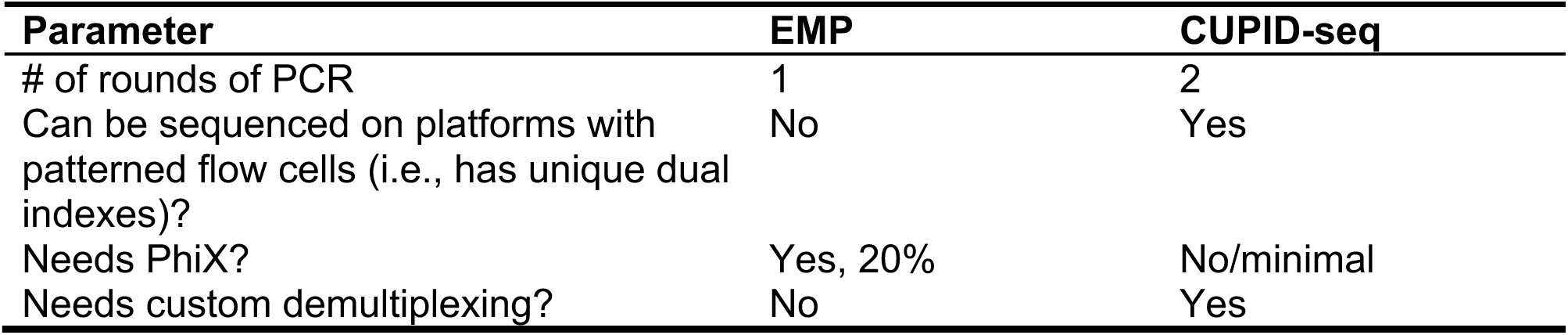
Pros and cons of the EMP protocol versus CUPID-seq.

| Parameter | EMP | CUPID-seq |
| --- | --- | --- |
| # of rounds of PCR | 1 | 2 |
| Can be sequenced on platforms with patterned flow cells (i.e., has unique dual indexes)? | No | Yes |
| Needs PhiX? | Yes, 20% | No/minimal |
| Needs custom demultiplexing? | No | Yes |

This combinatorial structure substantially increases multiplexing capacity (**Fig. 2**). For example, a NextSeq 2000 P1 600-cycle sequencing run produces ∼100 million reads, sufficient to sequence eight 96-well plates at an average depth of ∼120,000 reads per sample. Under conventional protocols, sequencing eight plates of samples simultaneously would require one Round 1 primer pair plus 8 × 96 Round 2 UDI primer pairs (1 + 768 = 769 primer pairs total). In contrast, CUPID-seq requires only 8 distinct Round 1 primer pairs (one per plate) and a single 96-well set of Round 2 UDI primers (8 + 96 = 104 primer pairs total), representing an ∼85% decrease in upfront primer requirements while preserving full sample distinguishability. In principle, even further reductions in the required number of primer pairs could be achieved by increasing the number of Round 1 primer pairs: for instance, by using 16 Round 1 primer pairs and 48 Round 2 UDI primer pairs to uniquely distinguish eight 96-well plates of samples.

**Figure 2:**
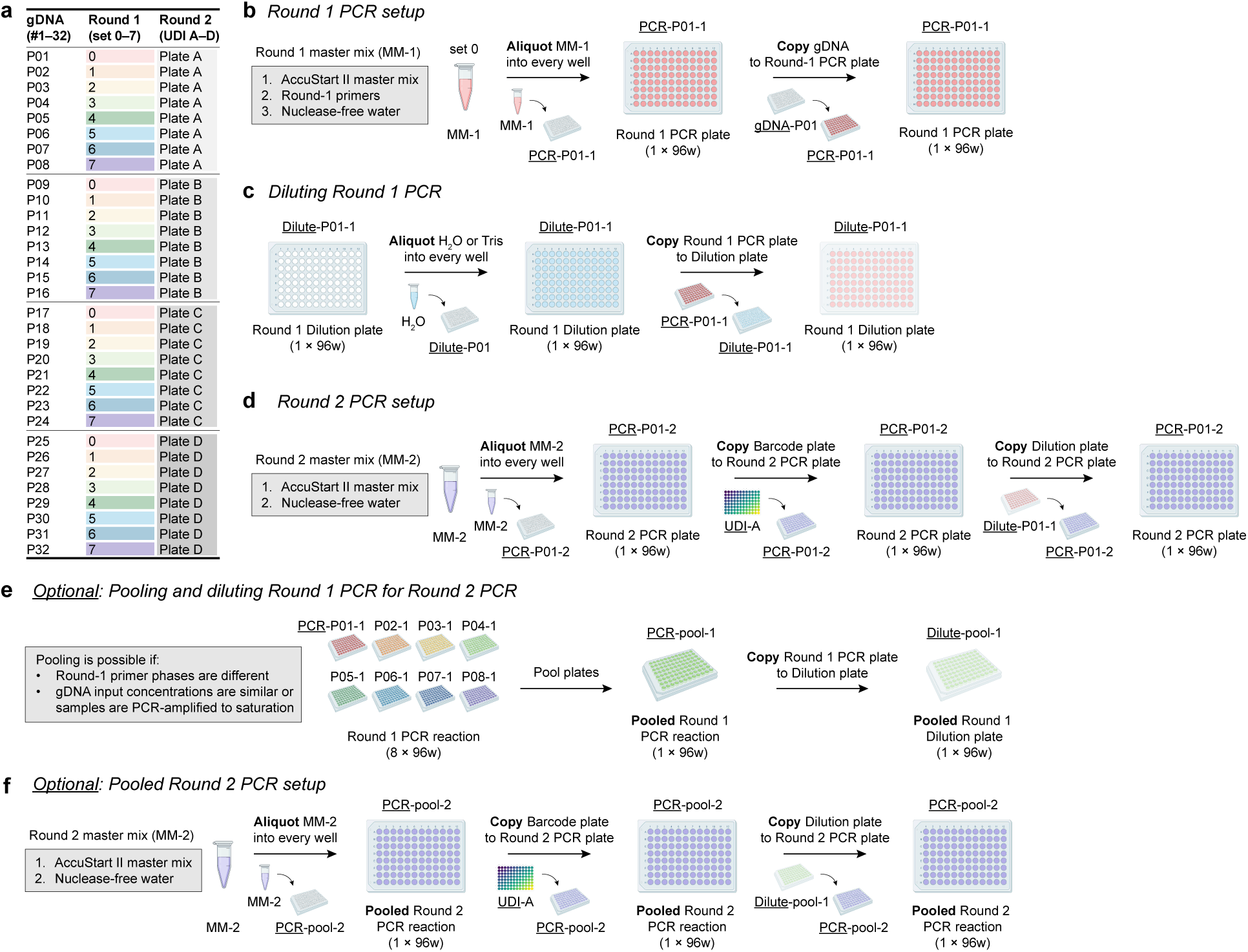
Schematic of experimental workflow for preparing 32 96-well plates. A) Mapping of unique Round 1 (primer sets 0–7) and Round 2 (barcode plates A–D) combinations to prepare 32 96-well plates for CUPID-seq. Panels B–D demonstrate how to prepare plates using Round 1 phase 0 primers and Round 2 UDI plate A. B) For Round 1 PCR, one master mix containing the appropriate Round 1 primer set should be prepared and aliquoted into a 96-well plate. Copy over the genomic DNA to each well. C) Following Round 1 PCR, samples should be diluted in either Tris buffer or water to decrease the concentration of Round 1 primers. D) For Round 2 PCR, the same master mix can be prepared and aliquoted into all 96-well plates. Copy over the appropriate Round 2 barcode plate, followed by the diluted Round 1 PCR samples. E) If samples are of similar DNA concentrations and utilize distinct Round 1 primer phases, up to 8 96-well plates may be pooled together and diluted. F) Pooled samples can be prepared in a single 96-well plate for Round 2 PCR.

In addition to reducing primer costs, CUPID-seq optionally decreases library preparation workload (**Table 3**). Conventional workflows require two PCRs per sample, resulting in 8 × 96 Round 1 reactions plus 8 × 96 Round 2 reactions (1536 PCRs total) to process eight plates. With CUPID-seq, up to eight Round 1 products bearing distinct in-line UDIs can be pooled prior to Round 2 amplification when input DNA concentrations are comparable or when Round 1 reactions are driven to saturation. In this configuration, eight plates can be prepared using 8 × 96 Round 1 reactions plus 96 Round 2 reactions (864 PCRs total), reducing reagent use and hands-on time by ∼40%. By distributing indexing across two PCR rounds, CUPID-seq leverages combinatorial scaling to align multiplexing capacity with modern sequencing throughput.

**Table 3:** Cost of preparing 8×96 samples for sequencing using the EMP protocol versus CUPID-seq. In this comparison, the EMP protocol processes 8 96-well plates (768 samples) sequenced on one MiSeq v3 600-cycle flow cell, while CUPID-seq processes 32 96-well plates (3,072 samples) on a single NovaSeq 6000 SP lane.

| Parameter | EMP | CUPID-seq |
| --- | --- | --- |
| DNA extraction | Same | Same |
| Round 1 PCR | \$50 per plate | \$50 per plate |
| Round 2 PCR | \$0 | \$50 per plate |
| Cost per sequencing run | \$2162 | \$2552.50 |
| Cost per M paired-end reads | \$86 | \$3 |
| Sequencing cost per 96-well plate | \$270 | \$80 |
| Coverage per sample | 10 K reads | >100 K reads |

### Overview of computational workflow

To complement the experimental workflow, we developed a computational pipeline to demultiplex CUPID-seq libraries and remove the Round 1 in-line UDIs from sequencing reads (**Fig. 1D**). We assume that sequencing data have already been demultiplexed based on Round 2 UDIs using standard Illumina software such as bcl2fastq and BCL Convert. However, no established tools are designed to demultiplex phased in-line indexes of variable length, motivating the development of a dedicated workflow.

The pipeline takes as input (i) the Round 1 index and primer sequences, (ii) a sample sheet linking Round 1 UDI and Round 2 UDI combinations to sample identifiers, (iii) a sheet linking fastq file paths to sample identifiers, and (iv) fastq files demultiplexed based on Round 2 UDIs (as performed by the sequencing provider or bcl2fastq/BCL Convert). Reads are first tagged according to their in-line Round 1 indexes using umi-tools^25^, enabling separation of reads into distinct index-defined groups. Residual index, spacer, and gene-specific primer sequences are then trimmed using cutadapt^26^, with parameters adjusted to accommodate variable index lengths. The resulting fastq files correspond to uniquely defined combinations of Round 1 and Round 2 UDIs.

Workflow execution and file management are coordinated using Snakemake^27^, and software dependencies (including Python^28^, Pandas^29^, NumPy^30^, SciPy^31^, Matplotlib^32^, in addition to cutadapt^26^ and umi-tools^25^) are encapsulated within Docker^33^ and Apptainer^34^ containers to ensure portability and reproducibility across local workstations and high-performance computing environments. Documentation and example datasets are provided to support straightforward implementation.

### Limitations of our approach

CUPID-seq optionally permits pooling of up to eight Round 1 PCR products bearing distinct in-line UDIs prior to Round 2 amplification (see **Anticipated results**, **Fig. 2**), reducing library preparation time and reagent use. However, pooled Round 2 PCRs may not be appropriate when samples have substantially different DNA concentrations or require different sequencing depths. In such cases, pooled amplification preserves differences in Round 1 product abundance and can result in increased variability in sequencing depth relative to independently amplified samples.

The use of phased in-line UDIs also increases computational complexity compared to constant-length in-line indexes, which are more straightforward to identify and trim. Although phased indexes introduce nucleotide diversity at the start of sequencing reads^23^ and reduce the requirement for PhiX DNA^24^, this feature may be unnecessary when base diversity is achieved through other means, such as co-sequencing shotgun libraries or partial-lane loading. In these situations, CUPID-seq can be simplified by using the provided 7-bp in-line index sequences without phasing (e.g., 3 bp on the forward primer and 4 bp on the reverse primer), although downstream demultiplexing will require custom modifications of the provided pipeline.

### Applications

CUPID-seq was developed and validated for amplification of the V4 region of the 16S rRNA gene, but the indexing architecture is readily adaptable to other 16S variable regions or user-defined amplicons by substituting the gene-specific primer regions (see “**Anticipated results**”, **Box 2**, **Table 4**). Because indexing is distributed across two PCR rounds, additional genomic targets can be incorporated without requiring a proportional expansion of Round 2 UDI primer sets.

**Table 4:**
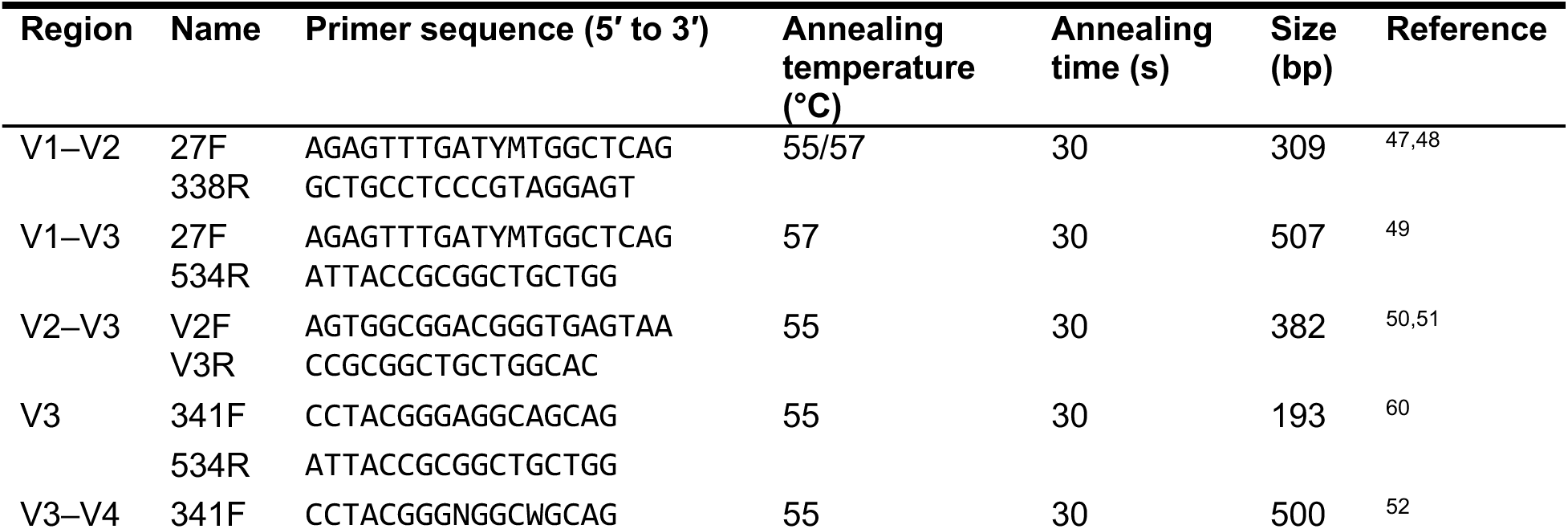
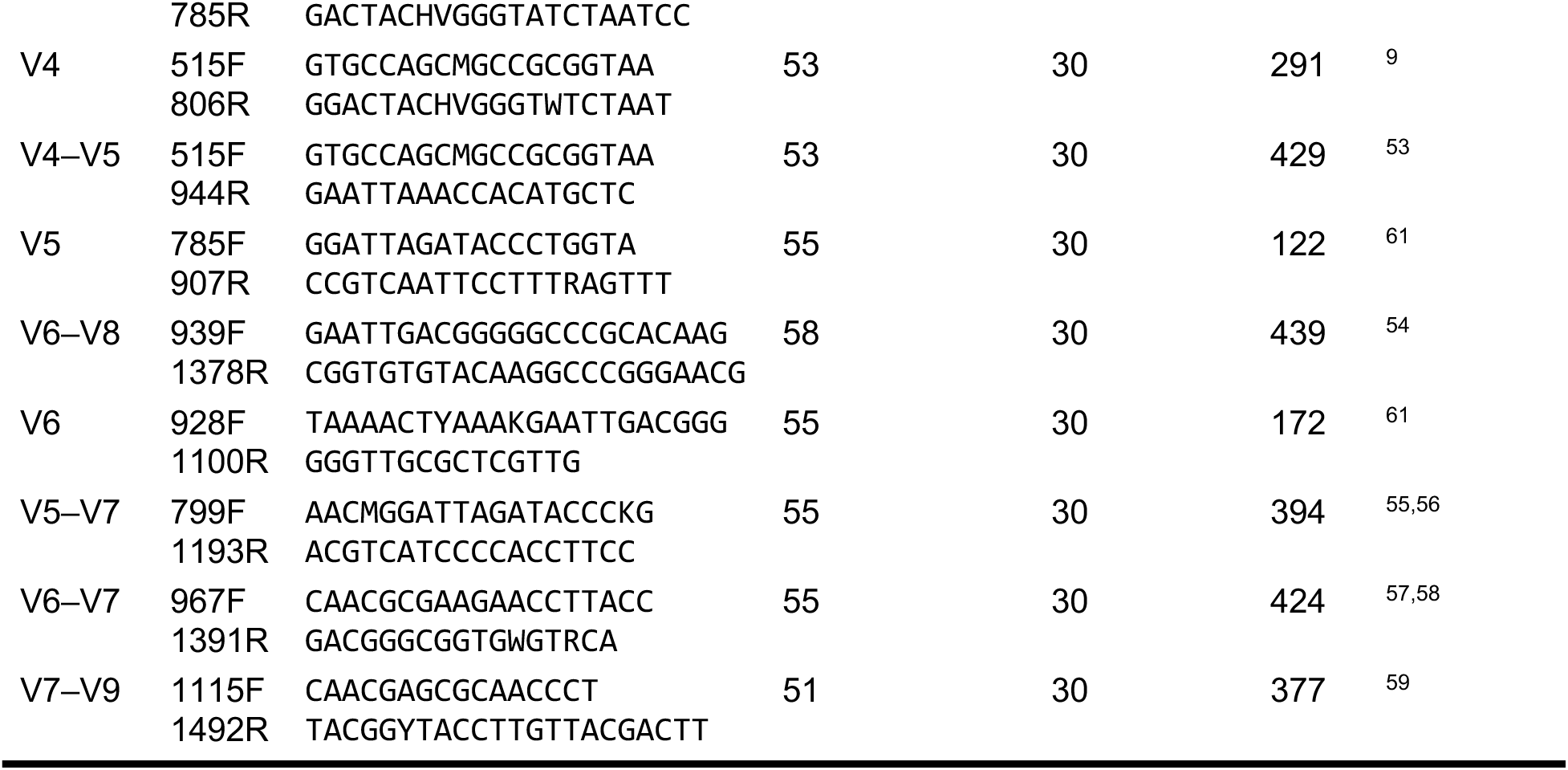
Primers for 16S rRNA variable regions.

This flexibility facilitates applications that require multiplexed profiling of multiple loci within the same study. For example, CUPID-seq can be adapted to detect mutations in multiple target genes^5^ or to concurrently profile bacterial and fungal^35^ communities alongside host-derived dietary markers^36^. By aligning indexing scalability with modern sequencing throughput, CUPID-seq extends high-capacity amplicon sequencing to experimental designs that were previously constrained by primer cost and multiplexing capacity.

### Experimental design

#### Designing Round 1 primers

Round 1 PCR is performed using up to eight primer pairs, each comprising the following elements (5′ to 3′):

##### Sequencing primer overhang

A constant sequence derived from the Illumina Nextera sequencing primer introduces homology for attachment of Round 2 PCR primers.

##### In-line index

Each primer pair contains one of eight phased in-line UDIs (0–7 bp total; **Table 5**). These indexes are positioned downstream of the gene-specific region and are sequenced at the beginning of each read. Distinct Round 1 in-line UDIs enable combinatorial barcoding when paired with Round 2 UDIs, allowing multiple samples to share the same Round 2 index set while remaining uniquely identifiable. The phased distribution of index lengths also introduces nucleotide diversity at the start of sequencing reads, reducing the need for PhiX DNA^24^.

**Table 5:** Phased in-line index sequences for Round 1 primers.

| Phase | Index F | Index R |
| --- | --- | --- |
| 0 | – | ATGGACT |
| 1 | T | GCTAGC |
| 2 | GG | TCACT |
| 3 | ACT | CGGT |
| 4 | TAAC | GTA |
| 5 | CAGTC | AA |
| 6 | ATCGAT | C |
| 7 | GCAAGTC | – |

The eight in-line UDIs were designed to satisfy the following criteria for V4 amplification:

1. The first seven sequenced bases differ by at least 3 bp between any pair of in-line UDIs (**Table 6**), reducing misassignment due to sequencing errors.
2. No single-nucleotide insertion or deletion converts one in-line UDI into another.
3. When all eight index sets are used simultaneously, each nucleotide (A, C, G, T) is represented at least once within the first 11 sequenced positions, ensuring that base diversity requirements are met^24^.

**Table 6:**
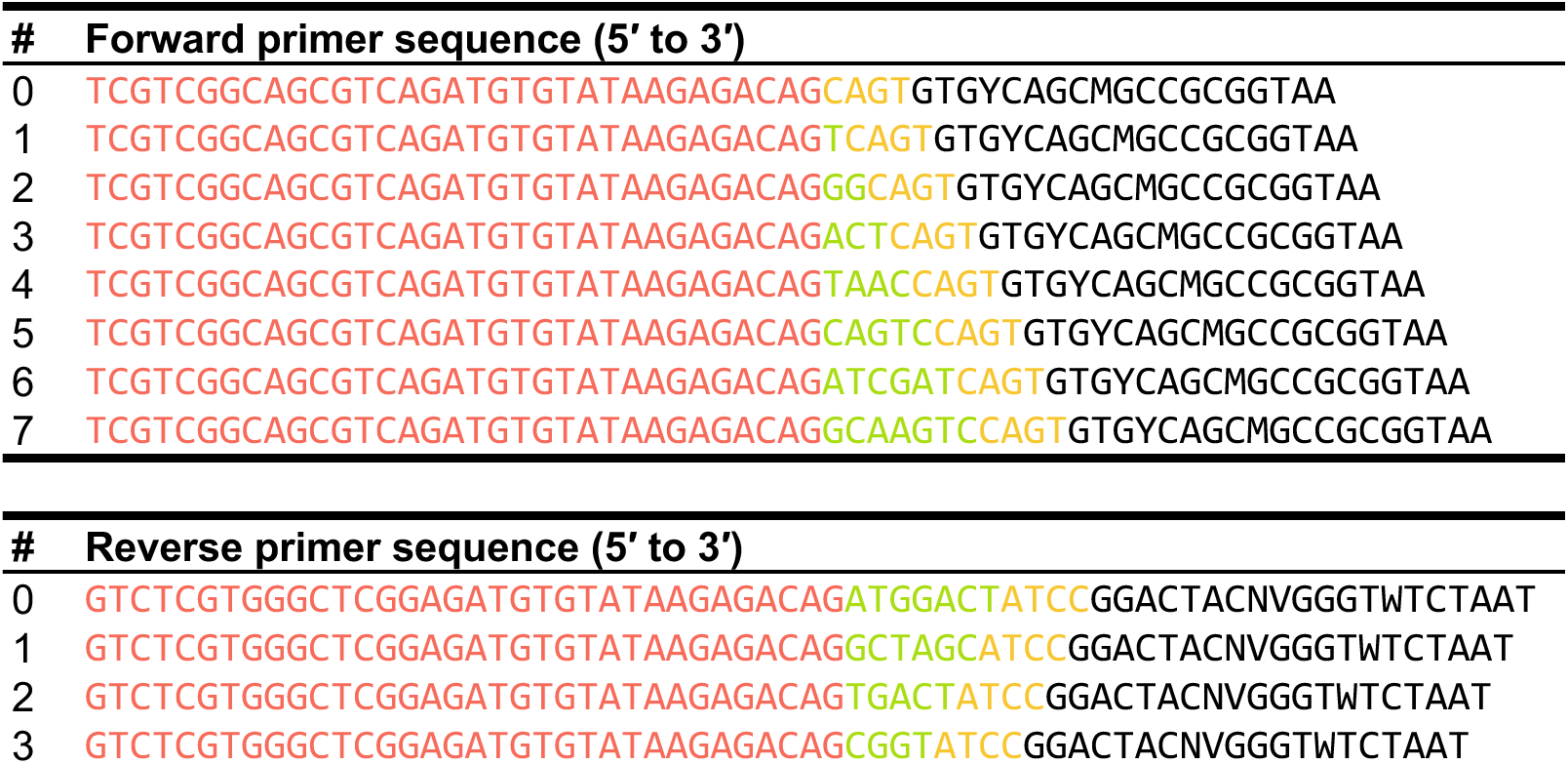
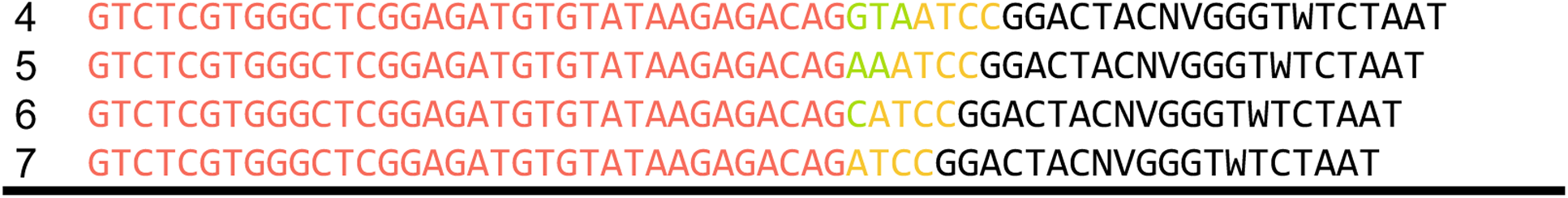
Round 1 primer sequences for amplification of the V4 region of the 16S rRNA gene. Each primer consists of a Round 2 homology region (red), in-line index (green), spacer region (yellow), and gene-specific sequence (black).

Scripts used to evaluate index design are available (checkBarcodes.R).

##### Spacer region

A constant 4-bp spacer is positioned between the gene-specific region and the in-line index to prevent unintended complementarity between the bases 5′ of the gene-specific sequence and the adjacent index, which could otherwise introduce amplification bias^9^. The two bases immediately adjacent to the gene-specific region (forward: GT; reverse: CC) match the pad sequences used in the EMP V4 primer set^9^. The remaining two bases were selected to increase nucleotide diversity during V4 sequencing.

#### Gene-specific region

These sequences determine target specificity and vary depending on the genomic locus of interest. For V4 amplification, Round 1 primers incorporate the 515F^37^ and 806R^38^ gene-specific sequences developed by the EMP^9^ (**Table 6**).

To adapt CUPID-seq to alternative targets, the gene-specific sequences in **Table 6** can be replaced with sequences specific to the region of interest. Annealing temperatures and other PCR conditions should be optimized accordingly (**Box 2**, **Table 4**). For convenience, Round 1 primers that target additional, commonly sequenced regions of the 16S rRNA gene are provided in **Table S1**, with gene-specific sequences and annealing temperatures provided in **Table 4**.

#### Round 1 PCR

Round 1 PCR amplifies the genomic region of interest from input DNA while incorporating the in-line UDI and a 5′ sequencing primer overhang that provides homology for Round 2 amplification. Each sample must be uniquely identifiable by a combination of its Round 1 and Round 2 UDI assignments within a given sequencing run. Accordingly, Round 1 PCR reactions are performed separately for each sample.

Annealing temperature, extension time, and number of cycles should be optimized for the target locus and input DNA quantity.

#### Round 2 PCR

Round 1 PCR products are diluted 1:40 in molecular-grade water to reduce residual primer carryover and used as template for Round 2 amplification^39^. When performing pooled Round 2 reactions, up to eight Round 1 products bearing distinct in-line UDIs can be combined prior to amplification (see “Anticipated results”). Pooled amplification is most effective when input DNA concentrations are normalized or when Round 1 reactions are driven to comparable endpoint amplification, minimizing variability in downstream sequencing depth (see “**Limitations of our approach**”).

Round 2 PCR is performed using Illumina Nextera Unique Dual Index primers. The standard Nextera UDI kit contains 384 pairs distributed across four 96-well plates (A–D). Representative primer sequences are shown in **Table 7**, and complete Illumina UDI sequences are available online (https://support-docs.illumina.com/SHARE/AdapterSequences/Content/Illumina-UDIndexes.htm). Adapter sequences required for sequencing center demultiplexing should be supplied at the time of submission.

**Table 7:**
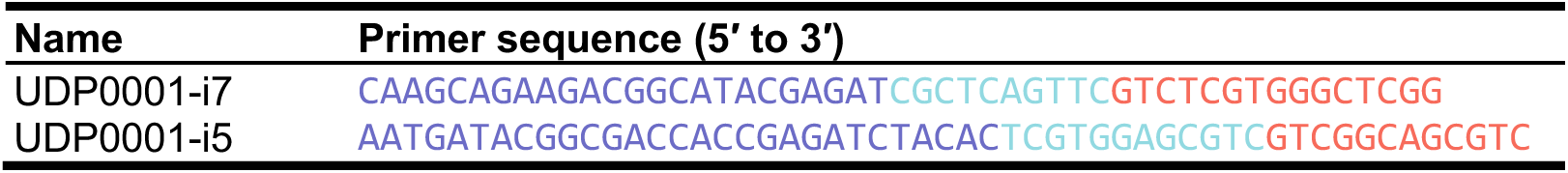
Example Round 2 primer sequences. Each primer comprises a flow-cell adapter (purple), unique dual index (blue), and a region homologous to the Round 1 PCR overhang (red).

Custom Round 2 primers incorporating alternative UDI sets may also be used. Round 2 primers should include the following regions (5′ to 3′):

##### Flow-cell adapters

Constant adapter sequences required for Illumina sequencing that enable library fragments to bind to the flow cell surface.

##### Index

UDI sequences incorporated during Round 2 PCR that permit sample demultiplexing and identification of index-hopping events.

##### Sequencing primer binding site

A region homologous to the Illumina Nextera sequencing primers that enables Round 2 primers to anneal to Round 1 PCR products.

#### Library cleanup and pooling

Prior to sequencing submission, Round 2 PCR products should be pooled and purified to remove residual primers and adapter dimers. Libraries may be combined either at equimolar concentrations or by volume, depending on the desired balance between workflow simplicity and evenness of sequencing depth.

For high-throughput applications, pooling by volume minimizes user time and reagent cost and has been previously validated^39^. In many cases, modest variability in sequencing depth can be addressed by sequencing to greater depth and subsampling reads during downstream analysis, offsetting the effort required to quantify and normalize the concentrations of hundreds of samples. However, when combining libraries derived from samples with substantially different input DNA concentrations (e.g., low-biomass vaginal samples and high-biomass stool samples), individual quantification and normalization are recommended to reduce depth variability.

Library purification can be performed using AMPure bead cleanups, PCR purification kits, or gel extractions. We provide a protocol for AMPure bead cleanup, which is also compatible with shotgun library preparation workflows, although alternative cleanup methods may be used as appropriate.

#### Demultiplexing and removal of Round 1 indexes

Sequencing providers typically demultiplex reads based on Round 2 UDIs. Additional demultiplexing based on Round 1 in-line UDIs is performed using the computational workflow described above. Reads are first separated according to Round 1 index assignments and subsequently trimmed to remove residual index, spacer, and gene-specific sequences.

Portable containerized environments are provided for both local workstations and virtual machines (Docker) and high-performance computing systems (Apptainer). These containers encapsulate nearly all required software dependencies, ensuring portability, reproducibility, and protection against version conflicts. The workflow supports both the primer sets described here and custom primer designs.

## Materials

### Biological samples

- Extracted genomic DNA

### Reagents

#### Chemicals

- Round 1 16S rRNA gene forward primers, salt-free (Integrated DNA Technologies, sequences provided in **Table 5**)
- Round 1 16S rRNA gene reverse primers, salt-free (Integrated DNA Technologies, sequences provided in **Table 5**)
- Nextera UDI Set A (Illumina, cat. no. 20091654)
- Nextera UDI Set B (Illumina, cat. no. 20091656)
- Nextera UDI Set C (Illumina, cat. no. 20091658)
- Nextera UDI Set D (Illumina, cat. no. 20091660)
- AMPure XP beads (Fisher Scientific, cat. no. A63880)
- AccuStart II PCR SuperMix (Quantabio, cat. no. 95137-04K)
- Ethanol, 200-proof, molecular grade (Millipore Sigma, cat. no. E7023-500ML)
- Elution buffer (10 mM Tris-HCl, pH 8.5) (Qiagen, cat. no. 19086)
- Molecular-grade water (Millipore Sigma, cat. no. 95284-100ML)
- Bovin serum albumin (Thermo Fisher Scientific, cat. no. B14)
- Dimethyl sulfoxide (Sigma–Aldrich, cat. no. D8418)
- Qubit dsDNA High Sensitivity Assay Kit (Thermo Fisher Scientific, cat. no. Q32851)

#### Plasticware and consumables

- 1,000-µL LTS filter pipette tips (Rainin, cat. no. 30389212)
- 200-µL LTS filter pipette tips (Rainin, cat. no. 30389239)
- 20-µL LTS filter pipette tips (Rainin, cat. no. 30389225)
- BenchSmart 1000-µL LTS filter pipette tips (Rainin, cat. no. 30296782)
- BenchSmart 200-µL LTS filter pipette tips (Rainin, cat. no. 17010646)
- BenchSmart 20-µL LTS filter pipette tips (Rainin, cat. no. 17011117)
- PCR strip tubes (Thermo Fisher Scientific, cat. no. AB-0264)
- Reservoirs (ChannelMate, cat. no. 1346-2510)
- DNA LoBind tubes, 1.5 mL (Eppendorf, cat. no. 022431021)
- 96-well PCR plates, skirted (Thermo Fisher Scientific, cat. no. AB-1400-L)
- 384-well PCR plates (Fisher Scientific, cat. no. 44-832-85)
- Aluminum seals for 96-well plates (Thermo Fisher Scientific, cat. no. 232698)
- Aluminum seals for 384-well plates (VWR, cat. no. 391-1281)

### Equipment

- BenchSmart 96 semi-automated pipetting system, 1000 µL (Mettler Toledo, cat. no. BST-96-1000) or equivalent
- BenchSmart 96 semi-automated pipetting system, 20 µL (Mettler Toledo, cat. no. BST-96-20) or equivalent
- Microcentrifuge (e.g., Thermo Fisher Scientific, cat. no. 75002436) or equivalent
- Plate roller (Andwin Scientific, cat. no. 60941118) or equivalent
- Mastercycler X50a PCR machine (Eppendorf, cat. no. 6313000018) or equivalent
- Pipet-Lite single-channel pipettes L-STARTXLS+ (Rainin, cat. no. 30456871) or equivalent
- Pipet-Lite multichannel pipette L12-200XLS+ (Rainin, cat. no. 17013810) or equivalent
- Pipet-Lite multichannel pipette L12-20XLS+ (Rainin, cat. no. 17013808) or equivalent
- Qubit 4 Fluorometer (Invitrogen, cat. no. Q33238) or equivalent

#### Hardware

- Desktop workstation or laptop with internet access (≥2 CPU cores and ≥64 GB of RAM recommended)
- Optional: high-performance computing (HPC) cluster with ≥64 GB RAM and Apptainer installed

#### Software

- Operating system: Linux, macOS, or Windows
- Python (version 3.13 or later)
- Pip (version 24.3 or later)
- CUPID-seq repository (http://github.com/KCHuang-Lab/CUPID-seq)
- Optional: Docker Engine (required for local or virtual machine execution, version 28.3 or later)
- Optional: Snakemake (required for HPC execution, version 9.3 or later)
- Optional: Apptainer (required for HPC execution, version 1.4 or later)
- Optional: snakemake-executor-plugin-slurm plugin (or equivalent; required for HPC execution via a resource manager)

#### Data files

Required input data:

- Demultiplexed raw sequencing reads (FASTQ format)
- fastq_list.txt, mapping forward and reverse reads to intermediate pooled sample identifiers
- samplesheet.txt, mapping pooled sample identifiers to final sample names and associated metadata
- Modified config.yaml file (included in repository)

Example datasets for testing the workflow are included within the CUPID-seq repository.

### Computational workflow setup

The computational workflow can be executed using either Docker (Procedure B Option 1; recommended for local or virtual machine execution) or Apptainer (Procedure B Option 2; recommended for HPC environments).

#### Procedure B Option 1 Setup: Docker installation and verification

To run the demultiplexing workflow locally, install and verify Docker Engine.

1. Install Docker Engine by downloading Docker Desktop from the official website (https://docs.docker.com/engine/install/) and following the installation instructions for the appropriate operating system.
2. Launch Docker Desktop.
3. Confirm that Docker is running by executing: docker version Successful installation and daemon execution will return version information for both the client and server components.

#### Procedure B Option 2 Setup: Apptainer setup and verification (HPC execution)

To run the workflow on an HPC cluster, Apptainer must be installed and accessible in the user environment.

1. Confirm that Apptainer is available on the HPC cluster by running: apptainer version If Apptainer is installed correctly, version information will be displayed.
2. If Apptainer is not installed, contact the HPC system administrator to request installation, or follow the official installation instructions provided at https://apptainer.org/docs/.
3. Install Snakemake according to the instructions provided here: https://snakemake.readthedocs.io/en/stable/getting_started/installation.html Confirm Snakemake is installed by running: snakemake -v If the installation was successful, the Snakemake version will be displayed.
4. Install the appropriate plugin for your cluster’s resource manager. For SLURM, the appropriate plugin can be installed by running: pip install snakemake-executor-plugin-slurm Additional plugins are described here: https://snakemake.github.io/snakemake-plugin-catalog/
5. Clone the CUPID-seq repository: git clone https://github.com/KCHuang-Lab/CUPID-seq.git If successful, the file structure shown in **Fig. S1** and **Fig. S2** should be created.

Note that Apptainer execution does not require root privileges. HPC resource allocation (including memory and CPU cores) should be specified according to cluster policies.

### Procedure A: Generating pooled libraries for amplicon sequencing

#### Procedure A1: Round 1 PCR

TIMING 30 min active and 2.5 h passive per 96-well plate of samples

CRITICAL Arrange samples such that each 96-well plate contains a single sample type (e.g., low-biomass saliva samples separate from high-biomass stool samples) to reduce variability in amplification efficiency and sequencing depth.

CRITICAL Each sample must have a unique combination of Round 1 UDI (Phase 0–7) and Round 2 UDI (plates A–D) within a sequencing run. A single Round 1 UDI can be used per plate if additional nucleotide diversity is provided by co-sequenced high-complexity (e.g., shotgun metagenomic) libraries. However, when sequencing amplicon libraries alone, use all eight Round 1 UDI sets for each Round 2 plate to maximize base diversity.

CRITICAL For low-biomass samples, perform PCR setup in a dedicated clean hood or PCR workstation to minimize environmental contamination.

1. Prepare Round 1 PCR reactions either individually, with the following mixture:

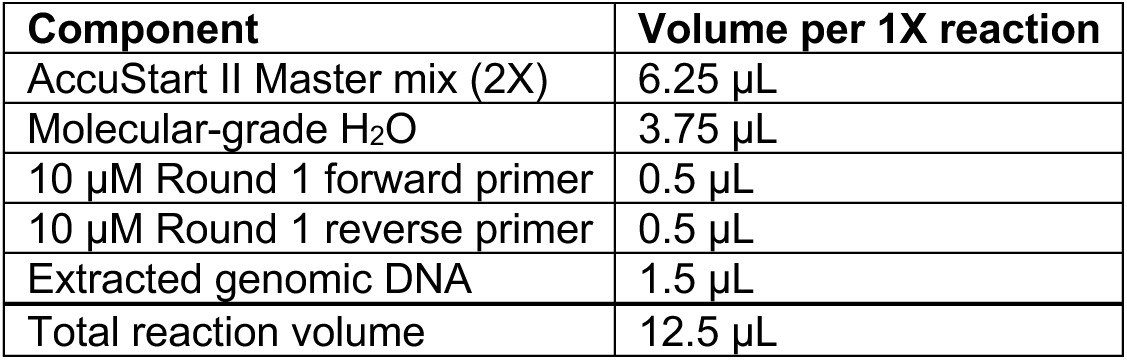

or as a master mix for one 96-well plate, using the following mixture (prepare ≥100 reactions to account for pipetting loss):

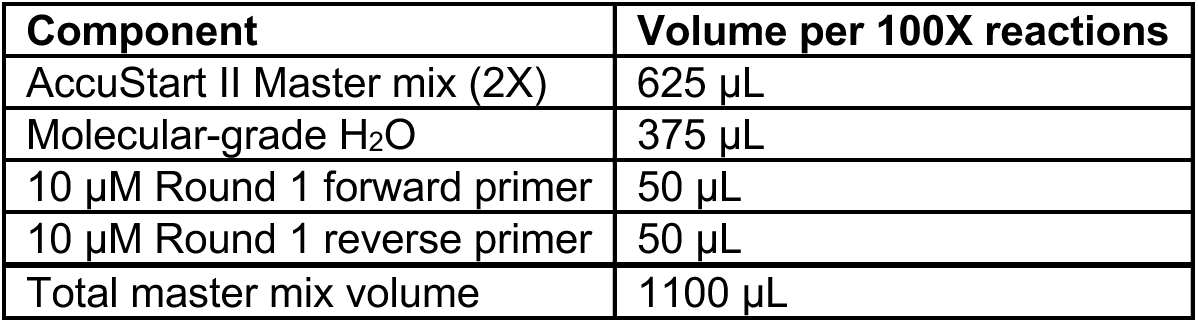 Aliquot 11 µL of master mix into each well. Add 1.5 µL of genomic DNA to each well to reach a final volume of 12.5 µL. CRITICAL Prepare a separate master mix for each Round 1 UDI set (Phase 0–7). CRITICAL High-biomass samples (e.g., stool) may contain PCR inhibitors. If amplification efficiency is reduced, supplement reactions with 0.5 µL of bovine serum albumin (BSA) or 0.5 µL of dimethyl sulfoxide (DMSO), and/or dilute genomic DNA prior to amplification.
2. Amplify the target locus using the following thermocycler conditions:

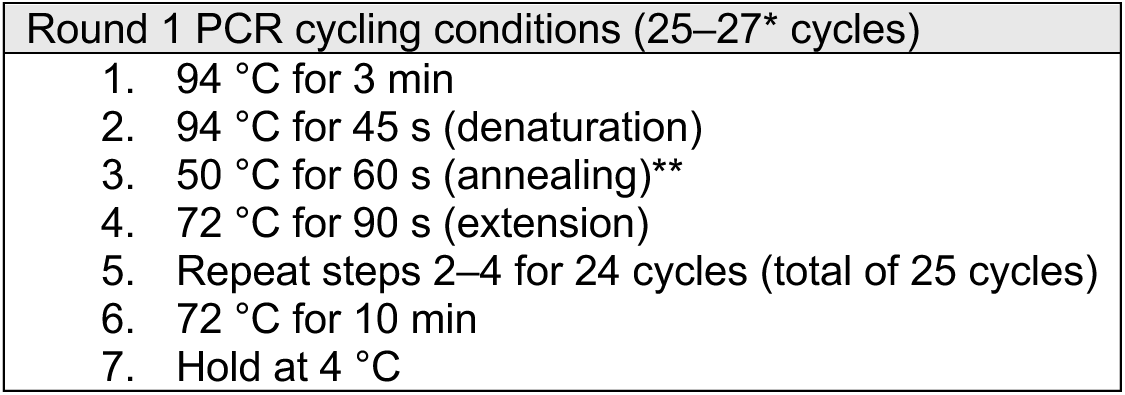 *: For cultivated communities and high-biomass samples (e.g., stool), 25–27 cycles are typically sufficient. For low-biomass samples, increase amplification to 30–35 cycles as needed. **: Annealing temperature should be optimized for each gene-specific region.
3. Verify amplification by running PCR products from at least one row per plate on a 1% (w/v) agarose gel prepared in TAE buffer. The expected amplicon size for 16S V4 amplification is 392 bp.

#### Procedure A2: Round 2 PCR

TIMING 1 h active and 1.5 h passive per 96-well plate of samples

1. Centrifuge Round 1 PCR plates and Round 2 primer plates at 3000*g* for 6 min to collect contents at the bottom of each well.
2. Dilute 2 µL of each Round 1 PCR product into 78 µL of 10 mM Tris buffer (pH 8.0) or nuclease-free H_2_O to dilute excess primers.

a. Optional: If Round 1 PCR products have similar DNA concentrations and desired sequencing depths, they can be pooled at the dilution step to decrease the number of Round 2 PCR reactions. Add equal volumes of each Round 1 PCR product into a new PCR plate and dilute 2 µL of the mixture of Round 1 PCR products into 78 µL of 10 mM Tris buffer (pH 8.0) or nuclease-free H_2_O. CRITICAL Pooled Round 1 samples must have distinct Round 1 UDIs. CRITICAL Do not use TE buffer for dilution, as EDTA will inhibit downstream reactions.
3. Prepare Round 2 PCR either individually with the following mixture:

| Component | Volume per 1X reaction |
| --- | --- |
| AccuStart II Master mix (2X) | 6.25 µL |
| Molecular-grade H<sub>2</sub>O | 3.25 µL |
| 5 µM Round 2 forward and reverse primer | 2 µL |
| Diluted Round 1 PCR product | 1 µL |
| Total volume | 12.5 µL |

or as a master mix for one 96-well plate format using the following mixture (prepare ≥100 reactions to account for pipetting loss):

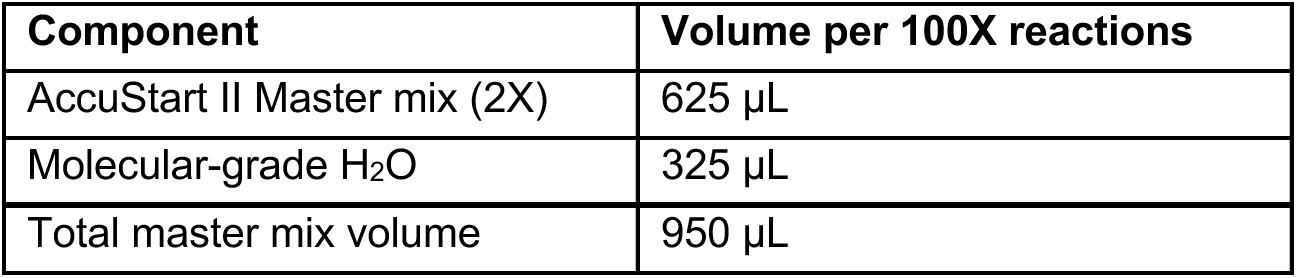 Aliquot 9.5 µL of master mix into each well. Add 1 µL of diluted Round 1 PCR product and 2 µL of the appropriate Round 2 UDI primer pair (5 µM each) to reach a total volume of 12.5 µL. CRITICAL Handle Round 2 UDI primer plates carefully to avoid cross-contamination. Reseal plates immediately after use and track usage to minimize primer degradation or well-to-well contamination. CRITICAL Use fresh filter tips when handling Round 2 primer plate stocks. Tips may be reused only when transferring between a Round 1 PCR product and corresponding dilution and Round 2 PCR plates, provided that a mixing step is included and no cross-sample transfer occurs.
4. Amplify the target locus using the following thermocycler conditions:

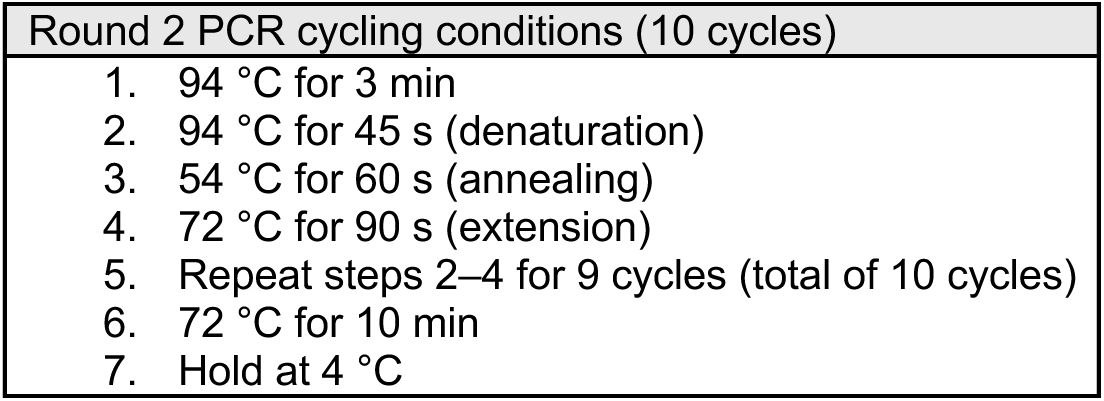 CRITICAL The annealing temperature for Round 2 PCR (54 °C) is higher than that used for Round 1 PCR due to the longer primer overhangs. This annealing temperature is determined by the Illumina primer sequence and does not need to be changed for different amplicon types.
5. Verify amplification by running PCR products from at least one row per plate on a 1% (w/v) agarose gel prepared in TAE buffer. The expected amplicon size for 16S V4 libraries after Round 2 PCR is 446 bp.

#### Procedure A3: Library cleanup and pooling

TIMING 1 h active per 96-well plate of samples

Prior to sequencing, Round 2 PCR products must be pooled and residual primers removed. Multiple pooling and cleanup strategies are compatible with this protocol. The procedure below describes pooling by equal volume followed by AMPure XP bead cleanup.

CRITICAL If multiple sample types are processed on the same plate and DNA concentrations vary substantially between wells, quantify DNA concentrations prior to pooling and combine samples at equimolar ratios.

CRITICAL Use low-DNA-binding tubes (e.g., LoBind) during this stage to minimize DNA loss.

CRITICAL Quantify DNA concentrations using a fluorometric assay (e.g., Qubit) rather than spectrophotometric measurement (e.g., Nanodrop) to ensure accurate library quantification.

1. Pool 2 µL of each Round 2 PCR product into a single tube (∼200 µL total per 96-well plate).
2. Perform a 0.8X AMPure XP bead cleanup of each pooled library according to the manufacturer’s instructions. Elute DNA in 50 µL of 10 mM Tris-HCl (pH 8.5).
3. Quantify the DNA concentration of each cleaned pool using a Qubit fluorometer.
4. If multiple plates are to be sequenced together, generate one pool per plate. To combine pools, dilute and normalize the volumes of each cleaned pool into a final sequencing library in nuclease-free water at the concentration requested by the sequencing provider.

### Procedure B: Demultiplexing samples and removing Round 1 indexes

The general file structure for the demultiplexing analysis is shown in **Fig. S1** and **Fig. S2**. Within the main folder 16S-demux, there are two important folders: config and workflow. Code for running the analysis is stored in workflow/rules/, and outputs are generated within workflow/out/. Test files (test_config.yaml, test_fastq_list.txt, and test_samplesheet.txt) and most input files are found within the config/ folder. The user-provided fastq_list.txt, and samplesheet.txt should also be included in the config folder. The names of these two files are flexible, but config.yaml assumes these defaults, and will need to be updated to reflect alternate names. Details on creating these input files are provided in **Box 1**.

The index_for_demux.txt file includes the in-line UDIs. By default, the 16S V4 indexes are used, so this file must be swapped to demultiplex other 16S regions or custom primers. Additional index_for_demux.txt files for other regions of the 16S rRNA gene are provided in the Github repository under other_index_for_demux.

The Snakefile documents encoding the analyses are found in the top-level 16S-demux folder.

CRITICAL Most common issues during demultiplexing arise from errors in the input files. To help with troubleshooting, the pipeline checks the input files before beginning operations and outputs a summary to 16S-demux/workflow/out/input_check_log.txt (or 16S-demux/workflow/test_out/input_check_log.txt for tests). This log may be helpful for troubleshooting errors that occur during the actual run but not in the test run.

#### Box 1

Inputs for the computational workflow

File formatting requirements and suggested conventions are provided below.

##### Sequencing data

- **Names:** Sample names should not include any spaces (underscores, periods, and hyphens are acceptable). If they do include such characters, the files should be renamed before demultiplexing.
- **Format:** Files should be formatted as gzipped fastq files (fastq.gz).
- **Location:** The raw sequencing fastq files must be accessible from wherever the demultiplexing analysis is run. The paths to fastq files are defined by the entries included in fastq_list.txt and the fastqdir variable in config.yaml. If the absolute path to the fastq files is provided in fastq_list.txt, then the fastqdir variable in config.yaml should be an empty string. Otherwise, ensure that the raw sequencing data can be accessed by concatenating the fastqdir path from config.yaml with the paths in fastq_list.txt. For example, the file /home/users/TEST/seq/TEST_R1_001.fastq.gz could be accurately described using an empty string within config.yaml and the full path /home/users/TEST/seq/TEST_R1_001.fastq.gz within fastq_list.txt. An alternative solution would be to set the fastqdir variable in config.yaml to /home/users/TEST/seq/ and to list the file name TEST_R1_001.fastq.gz in fastq_list.txt.

##### fastq_list.txt

- **Location:** fastq_list.txt should be located within the config directory, and the file name should be updated in the fastq_list field of the config.yaml file if not the default (fastq_list.txt).
- **Contents:** fastq_list.txt should be a table in a tab-delimited text file (.tsv or .txt). This table should have headers of read1, read2, and file, which refer to the path to read 1, the path to read 2, and the shortened filename, respectively. Additional columns will be ignored. For the filename, we recommend a format such as {run_name}-{R2_plate}-{well}, where run_name can be any string without underscores, spaces, or periods, and the next two terms specify the plate identifier for the Round 2 barcodes and the well, respectively. It is essential that the file field in the fastq file list matches the {run_name}-{R2_plate}-{well} portion of the filename field in samplesheet.txt. An example test_fastq_list.txt is shown in **Table 8**.

##### samplesheet.txt

- **Location:** samplesheet.txt should be included in the config directory, and the samplesheet path in the config.yaml file should be updated to reflect the samplesheet name if not the default (samplesheet.txt).
- **Contents:** The sample sheet should contain metadata for all samples as a tab-delimited table (.tsv or .txt). This file should contain a header as the first row, and must include the following columns: filename, sample, and group (**Table 9**). Additional columns can be included in the table as metadata for downstream analyses but will be ignored during sample demultiplexing. The filename column should have the format {run_name}-{R2_plate}-{well}-L{R1_index}, and the {run_name}-{R2_plate}-{well} portion should match the corresponding file entries in the fastq file list. The R1_index should match the phase entry in the indexfordemux.sh table. The sample column should contain the name assigned to each individual sample to take after demultiplexing and should not contain underscores or periods. The sample column value will be appended to the filename to form the final filename. Combinations of R1 and R2 UDIs that are not used in a sequencing run do not need to be included as rows in the sample sheet, even if the unused R1 UDIs are listed in config/index_for_demux.txt. If including these unused UDI combinations in the sample sheet, provide unique, non-NaN values for the sample and filename fields. The group column can contain any group identifier (without spaces or slashes). Samples in different groups will be output into different subdirectories within the trimmed directory at the end of the run. If reads do not need to be separated, use one group specifier for all samples, or leave the column blank while retaining the group header.

**Table 8:**
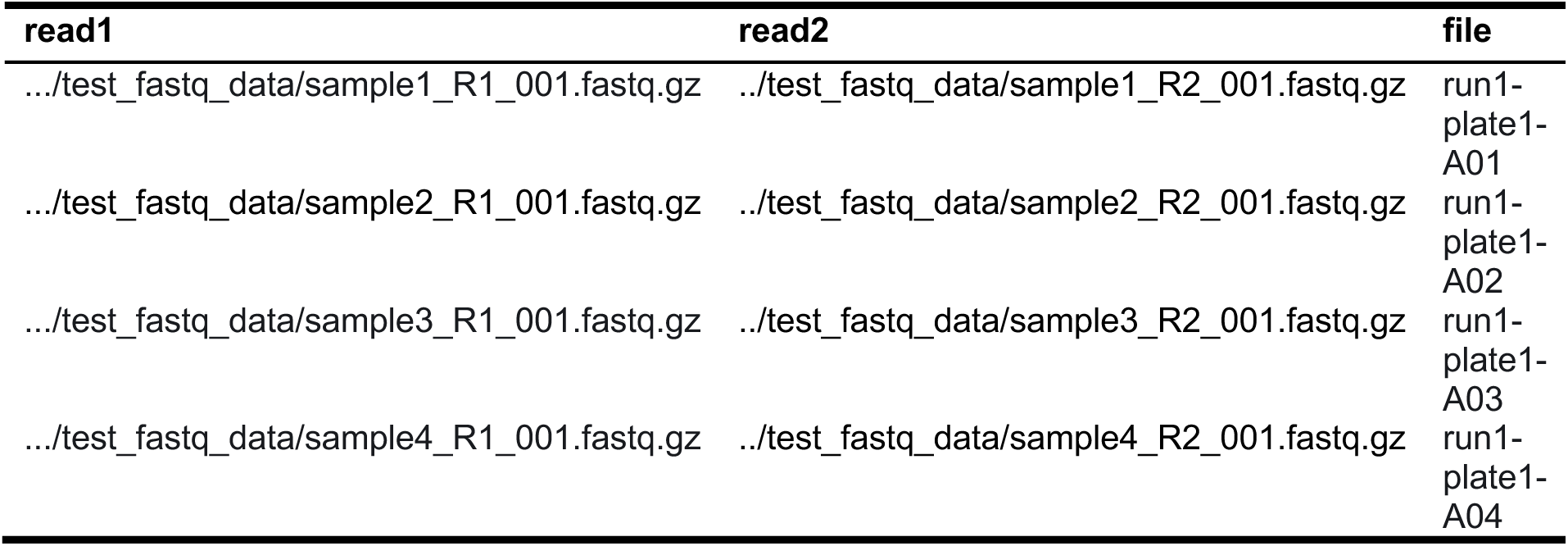
test_fastq_list.txt contents.

**Table 9:**
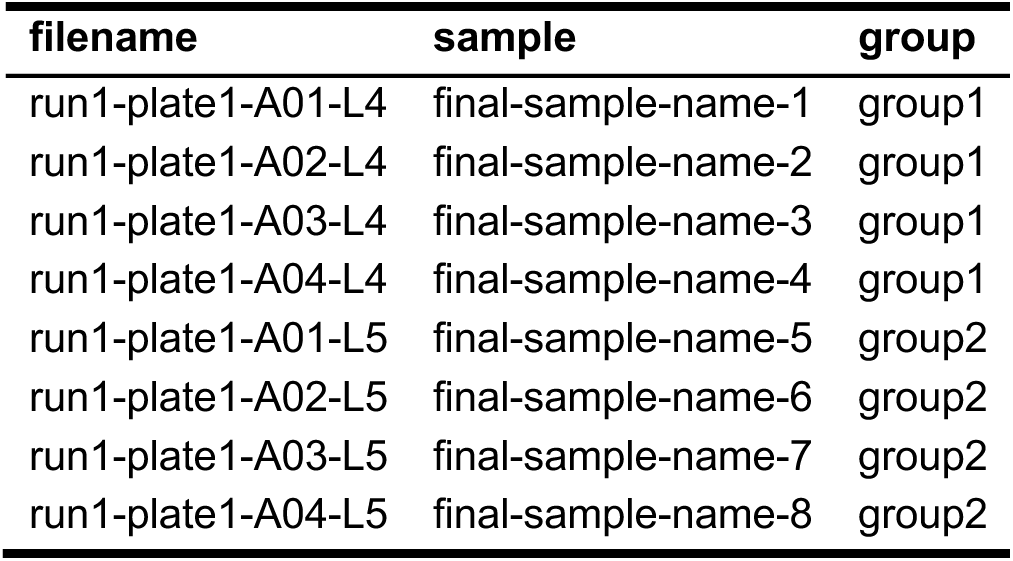
test_samplesheet.txt file contents.

#### Box 2

Integrating custom primers and indexes.

The instructions and code above assume that the primers and indexes used are the standard 16S V4 sets presented in the paper. If custom primers/indexes are used, several changes need to be made, as described below.

##### *Update* config/index_for_demux.txt

1. Make sure the index_for_demux.txt file, which is located in the config directory and specified in the config.yaml file, corresponds to the appropriate region. This file contains the index sequences used for demultiplexing R1 indexes. We provide primer sequences and corresponding index_for_demux.txt files for the following 16S regions: V1–V2, V1–V3, V2–V3, V3, V3–V4, V4–V5, V5, V5–V7, V6, V6–V7, V6–V8, and V7–V9. These files are provided in the other_index_for_demux folder of the GitHub repository (KCHuang-Lab/CUPID-seq), along with an Excel worksheet showing how these were derived and a template for generating sequences for additional amplicons (other_index_for_demux.xlsx).
2. If another region was amplified, the template in other_index_for_demux.xlsx can be used to make a custom index_for_demux.txt file. Replace the entries in geneF with the first part of the gene sequence (5′–3′, coding strand) and the entries in geneR with the end of the gene sequence (5′–3′, template strand) The Round 1 indexes have variable lengths as shown in **Table 5**. However, during demultiplexing, the reads are treated as if they have 7-bp indexes on both ends (**Table 10**). Any of the 7 bp that are not filled in with the index will be the spacer/gene-specific primer sequence. If the same primer design and index scheme are used, and only the gene-specific region is changed, then only the gene-specific regions within the 7-bp sequences need to be updated. For the primers in **Table 10**, the gene of interest starts with AGA… and ends with …GTA on the forward strand; therefore, the regions of homology include AGA and TAC, both in the 5′-to-3′ direction. For a gene reading ATG…CGT, the regions of homology within primers would become ATG and ACG, both in the 5′- to 3′- direction. Thus **index_for_demux.txt** should be edited (**Table 11**). CRITICAL We recommend avoiding mixed base characters such as W or N in the first three positions of the gene specific primer. If such bases are included, extra entries will need to be included in index_for_demux.txt, one for each potential base (e.g., for a W, one version should have an A and one should have a T). Each sample with a mixed base index should thus be included multiple times in samplesheet.txt, and the output files must then be merged downstream (either after trimming or after subsequent analyses).

##### Update the index lengths in config/config.yaml

Within the config/config.yaml file, the index/primer lengths may need to be updated. The lenR1index and lenR2index values should be set to the longest version of these indexes (e.g., in the provided primer/index set, the longest sequence of index bases is 7, so the value is set to 7). The lenR1primer and lenR2primer values should be set to equal the length of the gene-specific primer sequence plus the length of the spacer.

For the 16S primer sets with provided indexfordemux.txt files (V1–V2, V1–V3, V2–V3, V3, V3–V4, V4–V5, V5, V5–V7, V6, V6–V7, V6–V8, and V7–V9), the lengths do not need to be changed; the defaults in **Table 12** are correct.

**Table 10:** index_for_demux.txt file contents. The default indexes, with the index base pairs in red, the spacer base pairs in yellow, and the gene-specific primer regions highlighted in gray. The barcode column contains the concatenated strings of the two read indexes. The index, spacer, and gene-specific regions must be edited if sequencing other amplicons.

| Phase | Read_1_index | Read_2_index | Barcode |
| --- | --- | --- | --- |
| 0 | CAGTAGA | ATGGACT | CAGTAGAATGGACT |
| 1 | TCAGTAG | GCTAGCA | TCAGTAGGCTAGCA |
| 2 | GGCAGTA | TGACTAT | GGCAGTATGACTAT |
| 3 | ACTCAGT | CGGTATC | ACTCAGTCGGTATC |
| 4 | TAACCAG | GTAATCC | TAACCAGGTAATCC |
| 5 | CAGTCCA | AAATCCT | CAGTCCAAAATCCT |
| 6 | ATCGATC | CATCCTA | ATCGATCCATCCTA |
| 7 | GCAAGTG | ATCCTAC | GCAAGTGATCCTAC |

**Table 11:**
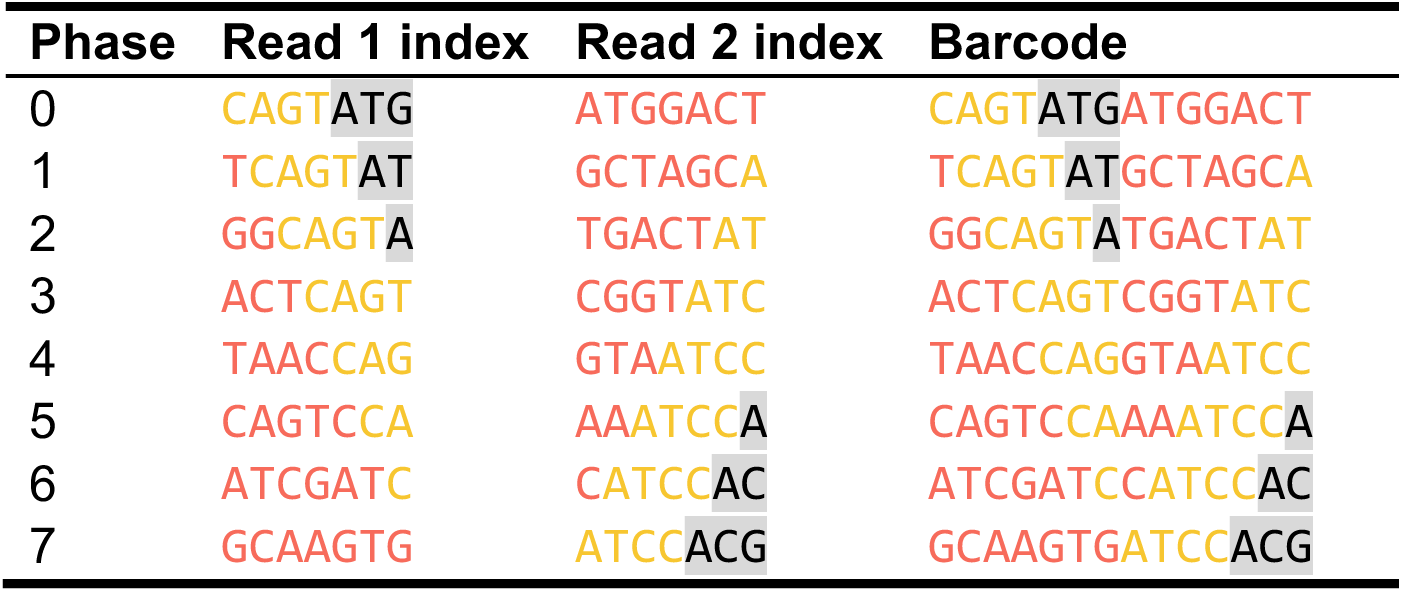
New index_for_demux.txt file contents.

**Table 12:** Length of indexes and primers.

| Region | Length |
| --- | --- |
| lenR1primer | 23 |
| lenR2primer | 24 |
| lenR1index | 7 |
| lenR2index | 7 |

### Procedure B Option 1: Using Docker

TIMING ∼2 hours

#### Installing and setting up the Docker image

The Docker image needed for the analysis can be acquired either by pulling directly from Docker Hub (the easiest approach) or by using the Dockerfile and other inputs provided in docker-demux.zip to build the Docker image.

##### (Sub-option 1) Pulling the image

TIMING ∼5 min

This option will make a local copy of the Docker image rlporter24/dualindex-demux, which contains the code and environment needed to process the data. Example input files for running a test are included.

1. Launch Docker.
2. In a terminal, run: docker pull rlporter24/dualindex-demux:1.0

##### (Sub-option 2) Building the Docker image

TIMING ∼10 min

The Docker image can also be built from the Dockerfile. (Note that this process takes longer and is not recommended.)

1. All files needed to build the Docker image are included in the GitHub repository at https://github.com/KCHuang-Lab/CUPID-seq. Clone the repository to create the file structure shown in **Fig. S1**. git clone https://github.com/KCHuang-Lab/CUPID-seq.git
2. Navigate to the demux directory: cd CUPID-seq/demux/ Within the docker_image_files directory are the Dockerfile and requirements.txt needed for the building process. The test_fastq_data directory contains example data for the test. All code files are contained within the 16S-demux directory, and output files are generated there as well.
3. From the demux directory, run the following: docker build -f docker_image_files/Dockerfile -t {name}:{version}. The image will be created locally with the name and version provided following -t. The build should take a few minutes and produce an output similar to this: => => naming to docker.io/library/demux/latest 0.0s View build details: docker-desktop://dashboard/build/docker-desktop-linux/argv3h81e8huy2fj9sown0 What’s next: View a summary of image vulnerabilities and recommendations -> docker scout quickview

#### Run the test analysis

TIMING ∼5 min

To check that the Docker image setup was successful, run an analysis using the provided test data. We recommend performing this test in interactive mode, as all commands can be run step by step within the container. All necessary input files are included within the container, so no files need to be imported.

1. To open a container from the image rlporter24/dualindex-demux in interactive mode (specified by the flag -it), run: docker run -it rlporter24/dualindex-demux:1.0 If a custom image was built, replace rlporter24/dualindex-demux:1.0 with the {name}:{version} provided for the build.
2. Once launched, the container should start in the 16S-demux directory containing both Snakefile (for the real analysis) and test_Snakefile (for the test analysis). If these files are not present within the current working directory, navigate to 16S-demux by running: cd /16S-demux
3. Run the test analysis, which uses the files config/test_fastq.txt, config/test_samplesheet.txt and test files included in /test_fastq_data/. with the command: snakemake --cores 1 -s test_Snakefile Using one core, this analysis should take under 5 minutes.
4. The output files will be generated in the workflow/test_out/ directory. If the run is successful, the outputs in **Fig. S3** and **Fig. S4** should be generated in workflow/test_out/trimmed.
5. Exit the container by running: exit Alternatively, stay in the container to run the actual analysis.

#### Run the actual analysis

TIMING ∼1 h for 384 input files with average size 7 GB, run with 4 cores and up to 10 GB memory.

1. If there is not already an open interactive container, launch one by running: docker run -it rlporter24/dualindex-demux:1.0 As before, if a custom image was built, replace rlporter24/dualindex-demux:1.0 with the {name}:{version} provided for the build.
2. Transfer the input files. The essential inputs will be the raw fastq data, fastq_list.txt, and samplesheet_txt. Details on creating these inputs for the actual analysis are included in **Box 1**. Inputs files can be transferred from inside or outside of the container (see https://docs.docker.com/reference/cli/docker/container/cp/). From outside the container (in a separate terminal), run: docker cp {local_path} {container_name}:{container_path} For example, run: docker cp ./fastqlist.txt optimistic_hawking:/16S-demux/config/ {local_path} can point to an individual file or a directory, and {container_name} should be the container name, not the image name. The container name can be found by running the command docker container ls to list all current containers, or by examining the containers in the Docker Desktop GUI. Docker container names generally follow the format of an adjective followed by an underscore and the surname of a notable scientist. The {container_path} can be provided relative to the 16S-demux directory, which is the home directory within the container.
3. In the config directory, update config.yaml so that samplesheet: and fastqlist: in lines 2 and 4 are followed by the paths to your input samplesheet and fastqlist files, respectively (**Box 1**). If using custom primers or indexes, the input file for indices or the lengths of read 1 and read 2 indexes and primers (lines 3, 5, 6, 7, 8) may need to be adjusted. For more details on custom primers, see **Box 2**.
4. Navigate to the 16S-demux directory where the Snakefile is located and submit the demultiplexing job by running snakemake --cores n where n is the desired number of cores. The analysis time will vary with the number and size of input files, as well as the machine used. If the run is successful, a message similar to the one below should be output: [Mon Jul 14 11:58:58 2025] Finished job 0. 30 of 30 steps (100%) done Complete log: .snakemake/log/2025-07-14T115513.645501.snakemake.log CRITICAL Before starting the actual run, the command snakemake -n can be used to perform a dry run. This test is helpful for ensuring that names and file locations are correct before starting the full run.

#### Transfer output files out

TIMING ∼5 min

1. Once the run is completed, output files must be transferred out of the container. To move files located in /16S-demux/workflow/out/ to a local destination, run the following: docker cp {container_name}:/16S-demux/workflow/out/ {local_path} For example, run: docker cp optimistic_hawking:/16S-demux/workflow/out/ ./results/

#### Clean up

TIMING ∼1 min

1. After the run is completed and outputs have been transferred, exit the container. This action will only remove the container that was built using this image, and not the image. Be sure any important data or intermediates are transferred out of the container before removing it. The remnants of the created container can be removed with the following command:

docker rm {container_name}

### Procedure B Option 2: Using Apptainer on a HPC cluster

TIMING ∼2 hours

High-Performance Computing (HPC) clusters that are not compatible with Docker may support Apptainer. Most HPC clusters use a resource manager, such as Simple Linux Utility for Resource Management (SLURM). The code includes example scripts for submitting through SLURM, but these scripts may need to be adapted for other clusters. Information regarding plugins for working with Snakemake using other job managers can be found here: https://snakemake.github.io/snakemake-plugin-catalog/.

#### Building the Apptainer image

TIMING ∼15 min

Ensure that Apptainer is installed. The pre-built Apptainer image is not currently available for direct download, but the Apptainer image can be built locally using files included in the GitHub repository. To build an Apptainer image, ensure that the necessary files described below are arranged properly, and build from the .def file.

All files needed to build the Apptainer image are included in the GitHub repository at https://github.com/KCHuang-Lab/CUPID-seq in demux/apptainer_image_files. 16S-demux.def and requirements.txt are both needed for the build process, and submit_build.sh is an example submission script for SLURM resource managers. The directory test_fastq_data contains example data for the test. All code files are contained within the 16S-demux directory, and output files will be generated there as well.

1. After cloning the repository, navigate to the CUPID-seq/demux/apptainer_image_files directory and execute the following command: apptainer build demux-image.sif 16S-demux.def The image will be created locally, and the file demux-image.sif will be created in the current working directory. The build should take less than 10 minutes and produce an output ending in a message similar to this: + rm -rf /apptainer_test/requirements.txt INFO: Adding runscript INFO: Creating SIF file… [=========================================================] 100 % 0s INFO: Build complete: demux-image.sif If the run is submitted via a resource manager, this output may be redirected to the output log. Once completed, the file demux-image.sif will be in the current working directory. This file, which encodes the Apptainer image, can be moved to a desired location, shared with other users on the same computing system, and reused for all CUPID-seq analyses. CRITICAL This step will likely require more resources than are available on a HPC login node, so use either a compute node or a job manager to allocate resources. A template SLURM script (submit_build.sh) is included in the apptainer-demux.zip file. Node/memory/time steps can be altered based on the cluster configuration.

#### Run the test analysis

TIMING ∼20 min

Running test analyses interactively is not recommended on HPC clusters that rely on job managers. Doing so may result in ‘out of memory’ errors that are difficult to debug. The testing instructions below describe how to initiate a test analysis using SLURM; the procedure should be adapted for other job managers.

1. To check that setup was successful, run an analysis using the provided test data by navigating to the 16S-demux/ directory and running the following command (or an analogous one for different resource managers): cd ../16S-demux/ snakemake --cores 1 -s test_Snakefile --use-singularity –executor slurm --profile config/profile/ If Snakemake is installed in a conda environment, ensure that the environment has been activated. Note that the exact syntax of a submission may vary with the Snakemake version used. The line above is compatible with version 9.16.3. This analysis should take ∼20 minutes depending on cluster usage and run parameters. As written, jobs will be submitted using the default parameters specified in the Snakemake profile 16S-demux/config/profile/config.yaml. It will be run using the config files config/test_fastq_list.txt and config/test_samplesheet.txt, as well as the test files included in /test_fastq_data/. If the run is successful, output files will be generated within the workflow/test_out/trimmed directory (**Fig. S4**). Resources allocated to any run are specified within the profile in config/profile/config.yaml. The allocated RAM (mem_mb), disk storage space (disk_mb), and runtime (runtime) can be edited here, which will likely be necessary for larger analyses. CRITICAL All files created or edited from within the container will be permanently edited within the local environment, so files do not need to be exported from the container to the local directory. However, users should be careful not to make any changes within the container that are not intended to be permanent (e.g., deleting files). CRITICAL To run a test analysis using an image file in a different location or with a different name, update the path to the image file within test_Snakefile by replacing “../apptainer_image_files/demux-image.sif” in line 4 with the correct path.

#### Run the actual analysis

TIMING ∼1 h for 384 input files with average size 7 GB, run with 4 cores and up to 10 GB memory.

As with the test analysis, we do not recommend running analyses interactively. The instructions below describe how to initiate a run using SLURM; the procedure should be adapted for other job managers.

1. Ensure that input files are defined properly and located in the correct directories (**Box 1**) and that the paths within config/config.yaml are correct.
2. Start a run with the command snakemake --cores 1 --executor slurm --use-singularity --profile config/profile/ If desired, edit the number of cores and the resource parameters within the profile (config/profile/config.yaml) based on system capacity to speed up the analysis as desired. CRITICAL All files created or edited from within the container will permanently change the local environment, so files do not need to be exported from the container. Users should be careful not to make any undesired changes within the container (e.g., deleting files). CRITICAL Before starting the actual run, the command snakemake -n can be used to perform a dry run. This test is helpful for ensuring that names and file locations are correct before starting the full run. CRITICAL To run the analysis using an image file in a different location or with a different name, the path within Snakefile must be updated to reflect this. Replace “../apptainer_image_files/demux-image.sif” in line 4 with the correct path.

#### Test run full outputs

Within the workflow directory, there should be a test_out subdirectory containing demux and trimmed subdirectories. In demux, there should be four sets of three files: each should have one .extract.log file, and two .fastq.gz files. There should also be two subdirectories: R1 and R2. The contents of R1 and R2 should have the same filenames that correspond to forward and reverse reads for the specified sample. Only the outputs for sample run1-plate1-A02 are shown in **Fig. S3**, but additional files corresponding to run1-plate1-A01, run1-plate1-A03, and run1-plate1-A04 should also be located within the demux directory. For each sample, there should be eight files ending with -L*.fastq.gz, where * is an integer from 0–7 (**Fig. S3**).

Within the trimmed directory, there should be two subdirectories: group1 and group2, each containing the subdirectories R1, R2, and removed, and two summary files: lowReadsSummary.txt and summary.txt (**Fig. S4**). R1 contains the forward reads, and R2 contains the reverse reads. These demultiplexed and trimmed samples can be used in any downstream analyses, such as inferring ASVs with DADA2^40^.

### Troubleshooting

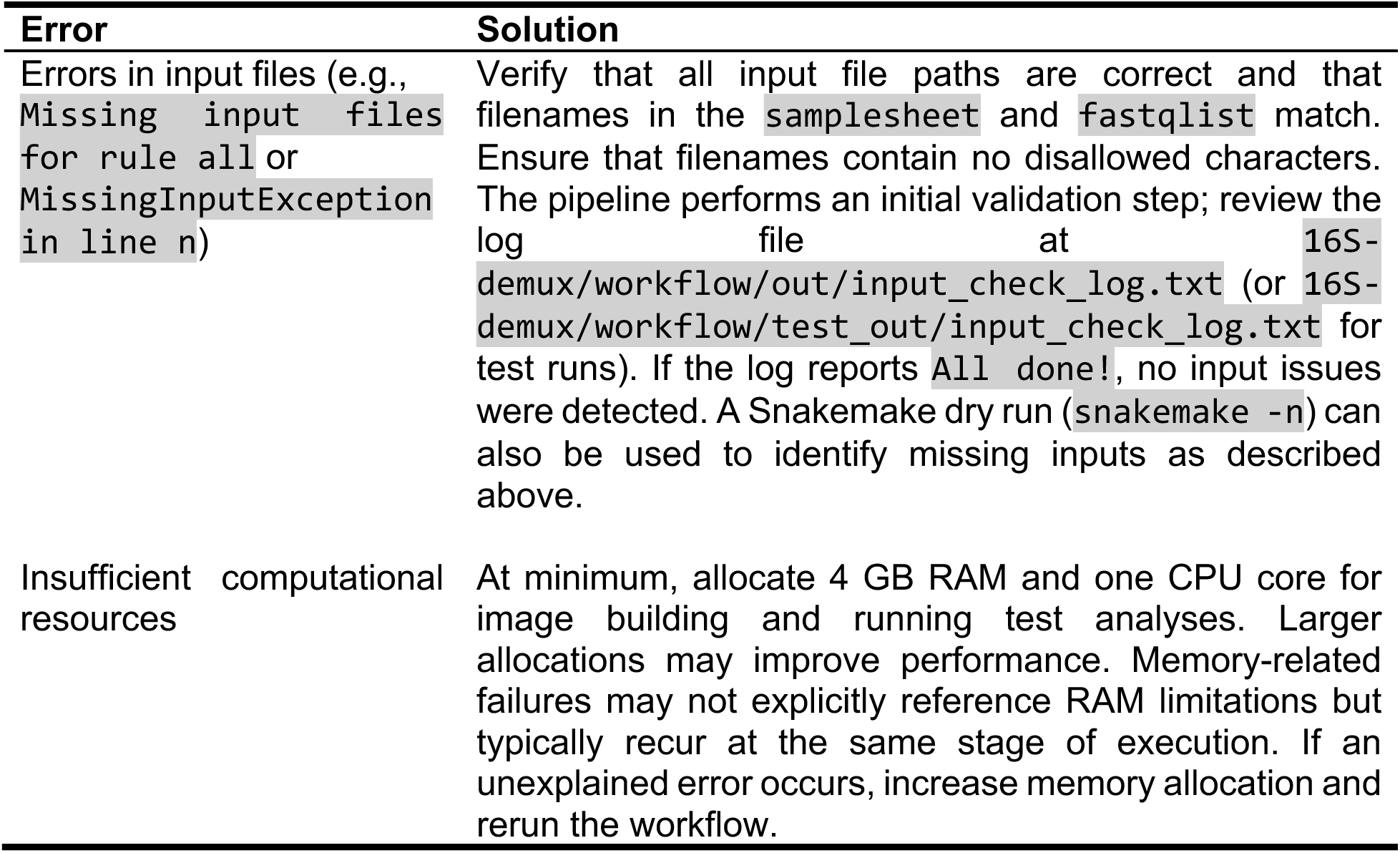

| Error | Solution |
| --- | --- |
| Errors in input files (e.g., Missing input files for rule all or MissingInputException in line n) | Verify that all input file paths are correct and that filenames in the samplesheet and fastqlist match. Ensure that filenames contain no disallowed characters. The pipeline performs an initial validation step; review the log file at 16S-demux/workflow/out/input_check_log.txt (or 16S-demux/workflow/test_out/input_check_log.txt for test runs). If the log reports All done!, no input issues were detected. A Snakemake dry run (snakemake -n) can also be used to identify missing inputs as described above. |
| Insufficient computational resources | At minimum, allocate 4 GB RAM and one CPU core for image building and running test analyses. Larger allocations may improve performance. Memory-related failures may not explicitly reference RAM limitations but typically recur at the same stage of execution. If an unexplained error occurs, increase memory allocation and rerun the workflow. |

### Timing

Procedure A1: Round 1 PCR: 30 min active and 2.5 h passive per 96-well plate of samples

Procedure A2: Round 2 PCR: 30 min active and 2.5 h passive per 96-well plate of samples

Procedure A3: Library cleanup and pooling: 1 h active per 96-well plate of samples

Procedure B: Demultiplexing samples and removing Round 1 indexes: ∼3 h for 384 samples, including initial setup (varies by sequencing depth and allocated resources)

### Anticipated results

#### Reproducibility across sequencing platforms, index sets, and PCR strategies

We evaluated the robustness of CUPID-seq to variation in sequencing platform, Round 1 in-line UDI choice, and pooled versus independent Round 2 PCR (**Fig. 3**). For validation, we amplified the V4 region of the 16S rRNA gene^9,41^ from genomic DNA extracted from strain isolates and complex *in vitro* microbial communities^39,42^ and inferred amplicon sequence variants (ASVs) using DADA2^40^.

**Figure 3:**
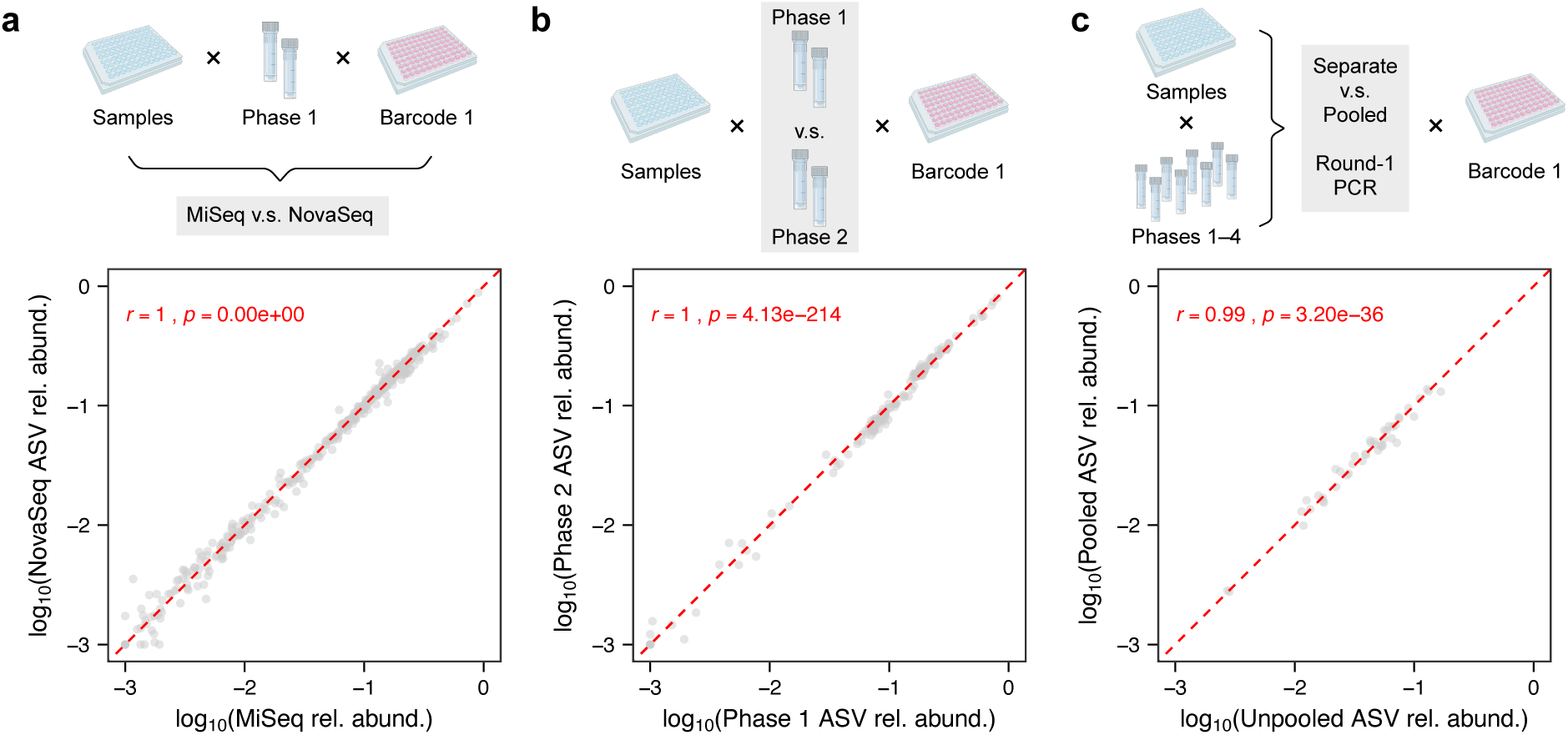
CUPID-seq is robust to the choice of sequencing platform, in-line index, and independent versus pooled Round 2 PCR. A) Sequencing of the same CUPID-seq library on MiSeq v3 (23 million clusters; non-patterned flow cells) and NovaSeq 6000 (800 million clusters; patterned flow cells) platforms resulted in highly correlated relative abundances for the most prevalent ASVs across all samples (*n* = 575). B) Sequencing of CUPID-seq libraries generated using distinct in-line (Round 1) indexes resulted in highly correlated relative abundances for the most prevalent ASV across all samples (*n* = 186). C) Using CUPID-seq, samples with unique in-line (Round 1) indexes can be pooled into a single Round 2 PCR without impacting community profiles. Sequencing of CUPID-seq libraries generated using independent versus pooled Round 2 PCRs resulted in highly correlated relative abundances for the most prevalent ASV across all samples (*n* = 44).

To assess platform effects, we prepared CUPID-seq libraries from six 96-well plates of *in vitro* communities and sequenced them on both the MiSeq v3 (non-patterned flow cell) and NovaSeq 6000 (patterned flow cell) platforms. Community composition was highly consistent between platforms, with strong correlation in the relative abundance of the most prevalent ASV (*Sutterella* sp.) in each sample between sequencing platforms (**Fig. 3A**; Pearson *r* = 1.00). Because ASV abundances are compositional and therefore interdependent, we focused on the most prevalent ASV in each sample to provide an interpretable metric of concordance. These results indicate that CUPID-seq libraries generated using different Illumina platforms can be directly compared^8^.

We next tested whether distinct Round 1 in-line UDIs introduced amplification or sequencing bias. Two 96-well plates were amplified using different Round 1 index sets and sequenced under identical conditions. Relative abundances of the most prevalent ASV (*Sutterella* sp.) were highly correlated between index sets (**Fig. 3B**; Pearson *r* = 1.00), indicating minimal bias attributable to in-line UDI choice. This robustness is consistent with the use of a constant 4-bp spacer between the gene-specific region and the index sequence.

Finally, we compared libraries generated using independent Round 2 PCRs with libraries prepared using pooled Round 2 amplification, in which multiple Round 1 products bearing distinct, in-line UDIs were combined prior to indexing. Relative abundances of the most prevalent ASV (*Flavonifractor* sp.) were strongly correlated between preparation strategies (**Fig. 3C**; Pearson *r* = 0.99), demonstrating that pooling does not introduce detectable amplification bias under the tested conditions. Furthermore, the dual-index architecture enables identification and exclusion of chimeric products generated during pooled Round 2 amplification^19^.

Collectively, these results demonstrate that CUPID-seq yields reproducible community composition profiles across sequencing platforms, Round 1 index sets, and optional pooled Round 2 amplification.

#### Comparison with the EMP protocol

Given the widespread use of 16S rRNA gene sequencing, we compared CUPID-seq with the EMP protocol, a commonly used method for amplification of the V4 region^8,9,37,38^ (**Table 2**, **Table 3**). The EMP protocol uses a single PCR to simultaneously amplify the target region and introduce a single index^9,37^. However, this design requires a uniquely barcoded forward primer per sample^37^, substantially increasing upfront primer costs for large studies—particularly on platforms capable of sequencing thousands of samples in a single run (**Table 1**). In addition, the absence of UDIs renders EMP libraries incompatible with patterned flow cell platforms due to index hopping^16,17^, and the protocol requires custom sequencing primers, complicating integration with other amplicon or shotgun libraries^43^. Although several two-step approaches address some of these limitations^23,44^, they do not incorporate dual rounds of unique indexing to enable combinatorial scaling.

To evaluate quantitative differences between protocols, we amplified identical samples using EMP primers and CUPID-seq primers containing the same gene-specific sequence to amplify the 16S V4 region (**Fig. 4A**). EMP libraries were sequenced on a MiSeq v3, as their single-index design is incompatible with patterned flow cell platforms, whereas CUPID-seq libraries were sequenced on a NovaSeq 6000 and subsampled to a comparable read depth. Based on our platform comparison (**Fig. 3A**), we anticipated minimal effects of sequencing platform on community composition. For additional context, we performed full-length 16S sequencing using PacBio for four *in vitro* communities (**Methods**).

**Figure 4:**
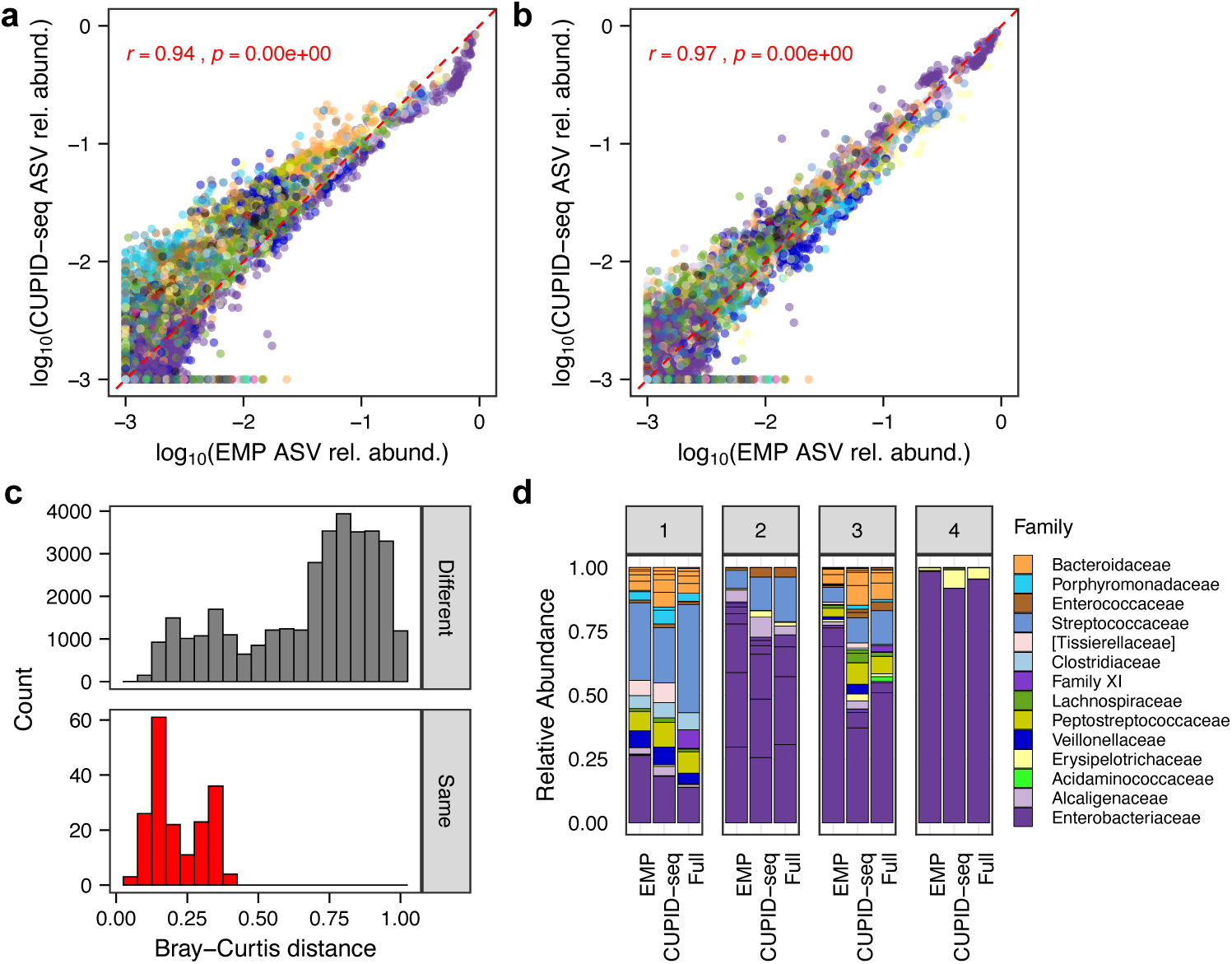
Comparison of EMP and CUPID-seq protocol amplification. A) The EMP and CUPID-seq protocols show systematic biases toward different bacterial families (*n* = 59,148 ASVs). For instance, CUPID-seq consistently estimates the relative abundance of Enterobacteriaceae ASVs (purple) at ∼70% of the value obtained with the EMP protocol, consistent with a ∼1.5% amplification bias per PCR cycle over 25 cycles. Each point represents a single ASV colored by its taxonomic family. B) Rescaling ASV abundances by the inferred family-level bias improves the correlation between the EMP and CUPID-seq protocols. Each point represents a single ASV colored by its taxonomic family. C) Comparison of Bray–Curtis distances between the same (bottom, red) or different (top, gray) samples. One sample in each pair was prepared using the EMP protocol, and the other using CUPID-seq. D) Family-level abundances of four *in vitro* communities using EMP (V4), CUPID-seq (V4), and full-length 16S sequencing. While there are quantitative differences among amplification methods, the variation across protocols is small compared to the biological differences across samples.

Across samples, relative abundances of ASVs were highly correlated between EMP and CUPID-seq datasets (Pearson *r* = 0.94). However, systematic fold-change differences were observed across taxonomic families, with median fold changes ranging from 0.71 to 4.28 (**Fig. 4A**, **Fig. S5**, **Table S2**). These shifts are consistent with modest amplification biases accumulated over 25 PCR cycles (up to ∼5% per cycle), within the range previously reported^39,45,46^.

To assess whether these systematic differences could be corrected, we calculated family-level fold changes and applied these correction factors to the data. After rescaling and normalization, agreement between protocols improved (Pearson *r* increased from 0.94 to 0.97; **Fig. 4A**, **Fig. 4B**, **Fig S6**, **Table S3**), indicating that amplification-related biases are predictable and partially correctable.

Although quantitative differences were detectable between EMP, CUPID-seq, and full-length 16S sequencing (**Fig. 4D**), inter-sample variation exceeded inter-protocol variation (**Fig. 4C**). These results reinforce prior observations^46^ that minor differences in primer architecture can introduce systematic shifts in relative abundance estimates, but that global community structure and beta-diversity patterns remain largely preserved across methods.

## Data and code availability

All data and code required for reproducing the analyses and figures are available on GitHub at https://github.com/fubeverly/CUPID-seq-paper. Code and documentation required for demultiplexing CUPID-seq libraries are available on GitHub at <u>https://</u> https://github.com/KCHuang-Lab/CUPID-seq. An archived version of the source files, including materials required to build Docker and Apptainer images from scratch, is deposited in the Stanford Digital Repository https://doi.org/10.25740/dt682kw9008. A pre-built Docker image for running the demultiplexing workflow is available on DockerHub (rlporter24/dualindex-demux:1.0).

## Supplemental Information

**Table S1:** Primer sequences and index_for_demux.txt parameters for amplifying different variable regions of the 16S rRNA gene.

**Table S2:** Pearson correlation coefficients (*r*) and FDR-corrected *p*-values for comparisons between EMP and CUPID-seq protocols at the family level.

**Table S3:** Pearson correlation coefficients (*r*) and FDR-corrected *p*-values for comparisons between EMP and CUPID-seq protocols at the family level after normalization. Relative abundances were normalized by the median fold change within each family.

## Author contributions statements

B.F., R.L.P., H.S., A.C.E., A.M.E., A.A, and K.S.X. carried out the experimental work and wrote the protocol. K.S.X. wrote the original manuscript and scripts. K.S.X., D.A.R, and K.C.H. conceived and supervised the original idea. All authors contributed to the review and editing of this manuscript and approved the final manuscript.

## Acknowledgments

The authors thank members of the Huang and Relman labs for helpful discussions. This work was supported by the National Science Foundation Postdoctoral Research Fellowship in Biology grant 2508176 (to B.F.), the National Science Foundation Graduate Research Fellowship Program award (to R.P.), National Institutes of Health grants T32 AI007328 (to B.F.), T32 GM136624 (to A.M.E.), R21 AI168860 (to D.R.), and DP1 DK147449 (to K.C.H.), a Department of Defense National Defense Science and Engineering Fellowship (to R.P.), the Thomas C. and Joan M. Merigan Endowment at Stanford University (to D.R.), James McDonnell Foundation Postdoctoral Fellowships in Understanding Dynamic and Multi-Scale Systems (to H.S. and K.S.X.), and the Jane Coffin Childs Memorial Fund Postdoctoral Fellowship (to K.S.X.). K.C.H. is a Chan Zuckerberg Biohub Investigator.

## References

1. van Opijnen, T., Bodi, K. L. & Camilli, A. Tn-seq: high-throughput parallel sequencing for fitness and genetic interaction studies in microorganisms. Nat. Methods 6, 767–772 (2009).

2. Wetmore, K. M. et al. Rapid Quantification of Mutant Fitness in Diverse Bacteria by Sequencing Randomly Bar-Coded Transposons. mBio 6, 10.1128/mbio.00306-15 (2015).

3. Fowler, D. M. & Fields, S. Deep mutational scanning: a new style of protein science. Nat. Methods 11, 801–807 (2014).

4. Levy, S. F. et al. Quantitative evolutionary dynamics using high-resolution lineage tracking. Nature 519, 181–186 (2015).

5. Chang, F. & Li, M. M. Clinical application of amplicon-based next-generation sequencing in cancer. Cancer Genet. 206, 413–419 (2013).

6. Clarridge, J. E. Impact of 16S rRNA Gene Sequence Analysis for Identification of Bacteria on Clinical Microbiology and Infectious Diseases. Clin. Microbiol. Rev. 17, 840–862 (2004).

7. Johnson, J. S. et al. Evaluation of 16S rRNA gene sequencing for species and strain-level microbiome analysis. Nat. Commun. 10, 5029 (2019).

8. Caporaso, J. G. et al. Ultra-high-throughput microbial community analysis on the Illumina HiSeq and MiSeq platforms. ISME J. 6, 1621–1624 (2012).

9. Caporaso, J. G. et al. Global patterns of 16S rRNA diversity at a depth of millions of sequences per sample. Proc. Natl. Acad. Sci. 108, 4516–4522 (2011).

10. Tringe, S. G. & Hugenholtz, P. A renaissance for the pioneering 16S rRNA gene. Curr. Opin. Microbiol. 11, 442–446 (2008).

11. Field, K. G. et al. Molecular Phylogeny of the Animal Kingdom. Science 239, 748–753 (1988).

12. Banos, S. et al. A comprehensive fungi-specific 18S rRNA gene sequence primer toolkit suited for diverse research issues and sequencing platforms. BMC Microbiol. 18, 190 (2018).

13. White, T. J., Bruns, T., Lee, S. & Taylor, J. Amplification and direct sequencing of fungal ribosomal RNA genes for phylogenetics. in PCR Protocols: A Guide to Methods and Applications 315–322 (Academic Press, Inc., 1990).

14. Walters, W., et al. Improved Bacterial 16S rRNA Gene (V4 and V4-5) and Fungal Internal Transcribed Spacer Marker Gene Primers for Microbial Community Surveys. mSystems 1, 10.1128/msystems.00009-15 (2015).

15. Taberlet, P., Coissac, E., Pompanon, F., Brochmann, C. & Willerslev, E. Towards next-generation biodiversity assessment using DNA metabarcoding. Mol. Ecol. 21, 2045–2050 (2012).

16. Sinha, R. et al. Index switching causes “spreading-of-signal” among multiplexed samples in Illumina HiSeq 4000 DNA sequencing. 125724 Preprint at 10.1101/125724 (2017).

17. Costello, M. et al. Characterization and remediation of sample index swaps by non-redundant dual indexing on massively parallel sequencing platforms. BMC Genomics 19, 332 (2018).

18. Shen, M.-J. R., et al. Kinetic exclusion amplification of nucleic acid libraries. (2013).

19. Kircher, M., Sawyer, S. & Meyer, M. Double indexing overcomes inaccuracies in multiplex sequencing on the Illumina platform. Nucleic Acids Res. 40, e3 (2012).

20. Effects of Index Misassignment on Multiplexing and Downstream Analysis. https://www.illumina.com/content/dam/illumina-marketing/documents/products/whitepapers/index-hopping-white-paper-770-2017-004.pdf?linkId=36607862.

21. Understanding unique dual indexes (UDI) and associated library prep kits. https://knowledge.illumina.com/library-preparation/general/library-preparation-general-reference_material-list/000002344 (2025).

22. 16S Metagenomic Sequencing Library Preparation. https://support.illumina.com/documents/documentation/chemistry_documentation/16s/16s-metagenomic-library-prep-guide-15044223-b.pdf.

23. Holm, J. B. et al. Ultrahigh-Throughput Multiplexing and Sequencing of >500-Base-Pair Amplicon Regions on the Illumina HiSeq 2500 Platform. mSystems 4, 10.1128/msystems.00029-19 (2019).

24. Nucleotide Diversity. https://support-docs.illumina.com/SHARE/ClusterOptimize/Content/SHARE/ClusterOptimize/NucleotideDiversity.htm.

25. Smith, T., Heger, A. & Sudbery, I. UMI-tools: modeling sequencing errors in Unique Molecular Identifiers to improve quantification accuracy. Genome Res. 27, 491–499 (2017).

26. Martin, M. Cutadapt removes adapter sequences from high-throughput sequencing reads. EMBnet.journal 17, 10 (2011).

27. Mölder, F., et al. Sustainable data analysis with Snakemake. Preprint at 10.12688/f1000research.29032.1 (2021).

28. The Python Language Reference. Python documentation https://docs.python.org/3/reference/index.html.

29. McKinney, W. Data Structures for Statistical Computing in Python. SciPy 2010 https://doi.org/10.25080/Majora-92bf1922-00a (2010) doi:10.25080/Majora-92bf1922-00a.

30. Harris, C. R. et al. Array programming with NumPy. Nature 585, 357–362 (2020).

31. Virtanen, P. et al. SciPy 1.0: fundamental algorithms for scientific computing in Python. Nat. Methods 17, 261–272 (2020).

32. Hunter, J. D. Matplotlib: A 2D Graphics Environment. Comput. Sci. Eng. 9, 90–95 (2007).

33. Merkel, D. Docker: lightweight Linux containers for consistent development and deployment. Linux J 2014, 2:2 (2014).

34. Kurtzer, G. M., Sochat, V. & Bauer, M. W. Singularity: Scientific containers for mobility of compute. PLOS ONE 12, e0177459 (2017).

35. Schoch, C. L. et al. Nuclear ribosomal internal transcribed spacer (ITS) region as a universal DNA barcode marker for Fungi. Proc. Natl. Acad. Sci. 109, 6241–6246 (2012).

36. Superdock, D. K., Petrone, B. L., Kirtley, M. C. & David, L. A. From stool to sequence: decoding the human diet with FoodSeq. mSystems 10, e00158–25 (2025).

37. Parada, A. E., Needham, D. M. & Fuhrman, J. A. Every base matters: assessing small subunit rRNA primers for marine microbiomes with mock communities, time series and global field samples. Environ. Microbiol. 18, 1403–1414 (2016).

38. Apprill, A., McNally, S., Parsons, R. & Weber, L. Minor revision to V4 region SSU rRNA 806R gene primer greatly increases detection of SAR11 bacterioplankton. Aquat. Microb. Ecol. 75, 129–137 (2015).

39. Celis, A. I. et al. Optimization of the 16S rRNA sequencing analysis pipeline for studying in vitro communities of gut commensals. iScience 103907 (2022) doi:10.1016/J.ISCI.2022.103907.

40. Callahan, B. J. et al. DADA2: High-resolution sample inference from Illumina amplicon data. Nat. Methods 13, 581–583 (2016).

41. Thompson, L. R. et al. A communal catalogue reveals Earth’s multiscale microbial diversity. Nature 551, 457–463 (2017).

42. Aranda-Díaz, A. et al. Establishment and characterization of stable, diverse, fecal-derived in vitro microbial communities that model the intestinal microbiota. Cell Host Microbe 30, 260–272.e5 (2022).

43. Spiking custom primers into the Illumina sequencing primers | Illumina Knowledge. https://knowledge.illumina.com/library-preparation/general/library-preparation-general-reference_material-list/000001542 (2026).

44. Shahi, S. K., Zarei, K., Guseva, N. V. & Mangalam, A. K. Microbiota Analysis Using Two-step PCR and Next-generation 16S rRNA Gene Sequencing. J. Vis. Exp. JoVE https://doi.org/10.3791/59980 (2019) doi:10.3791/59980.

45. Hansen, M. C., Tolker-Nielsen, T., Givskov, M. & Molin, S. Biased 16S rDNA PCR amplification caused by interference from DNA flanking the template region. FEMS Microbiol. Ecol. 26, 141–149 (1998).

46. Polz, M. F. & Cavanaugh, C. M. Bias in Template-to-Product Ratios in Multitemplate PCR. Appl. Environ. Microbiol. 64, 3724–3730 (1998).

47. Salter, S. J. et al. Reagent and laboratory contamination can critically impact sequence-based microbiome analyses. BMC Biol. 12, 87 (2014).

48. Mesa, V. et al. Bacterial, Archaeal, and Eukaryotic Diversity across Distinct Microhabitats in an Acid Mine Drainage. Front. Microbiol. 8, (2017).

49. Walker, A. W. et al. 16S rRNA gene-based profiling of the human infant gut microbiota is strongly influenced by sample processing and PCR primer choice. Microbiome 3, 26 (2015).

50. Will, C. et al. Horizon-Specific Bacterial Community Composition of German Grassland Soils, as Revealed by Pyrosequencing-Based Analysis of 16S rRNA Genes. Appl. Environ. Microbiol. 76, 6751–6759 (2010).

51. Sahm, K. et al. High abundance of heterotrophic prokaryotes in hydrothermal springs of the Azores as revealed by a network of 16S rRNA gene-based methods. Extremophiles 17, 649–662 (2013).

52. Klindworth, A. et al. Evaluation of general 16S ribosomal RNA gene PCR primers for classical and next-generation sequencing-based diversity studies. Nucleic Acids Res. 41, e1 (2013).

53. Fuks, G. et al. Combining 16S rRNA gene variable regions enables high-resolution microbial community profiling. Microbiome 6, 17 (2018).

54. Lebuhn, M. et al. Towards molecular biomarkers for biogas production from lignocellulose-rich substrates. Anaerobe 29, 10–21 (2014).

55. Chelius, M. K. & Triplett, E. W. The Diversity of Archaea and Bacteria in Association with the Roots of Zea mays L. Microb. Ecol. 41, 252–263 (2001).

56. Bodenhausen, N., Horton, M. W. & Bergelson, J. Bacterial communities associated with the leaves and the roots of Arabidopsis thaliana. PloS One 8, e56329 (2013).

57. Sogin, M. L. et al. Microbial diversity in the deep sea and the underexplored “rare biosphere”. Proc. Natl. Acad. Sci. 103, 12115–12120 (2006).

58. Walker, J. J. & Pace, N. R. Phylogenetic Composition of Rocky Mountain Endolithic Microbial Ecosystems. Appl. Environ. Microbiol. 73, 3497–3504 (2007).

59. Turner, S., Pryer, K. M., Miao, V. P. W. & Palmer, J. D. Investigating Deep Phylogenetic Relationships among Cyanobacteria and Plastids by Small Subunit rRNA Sequence Analysis. J. Eukaryot. Microbiol. 46, 327–338 (1999).

60. Muyzer, G., De Waal, E. C. & Uitterlinden, A. G. Profiling of complex microbial populations by denaturing gradient gel electrophoresis analysis of polymerase chain reaction-amplified genes coding for 16S rRNA. Appl. Environ. Microbiol. 59, 695–700 (1993).

61. 16S rRNA and 16S rRNA Gene – EzBioCloud Help center. https://help.ezbiocloud.net/16s-rrna-and-16s-rrna-gene/.

