## Supplemental Information for "CUPID-seq enables highly multiplexed amplicon sequencing via combinatorial in-line dual indexing"

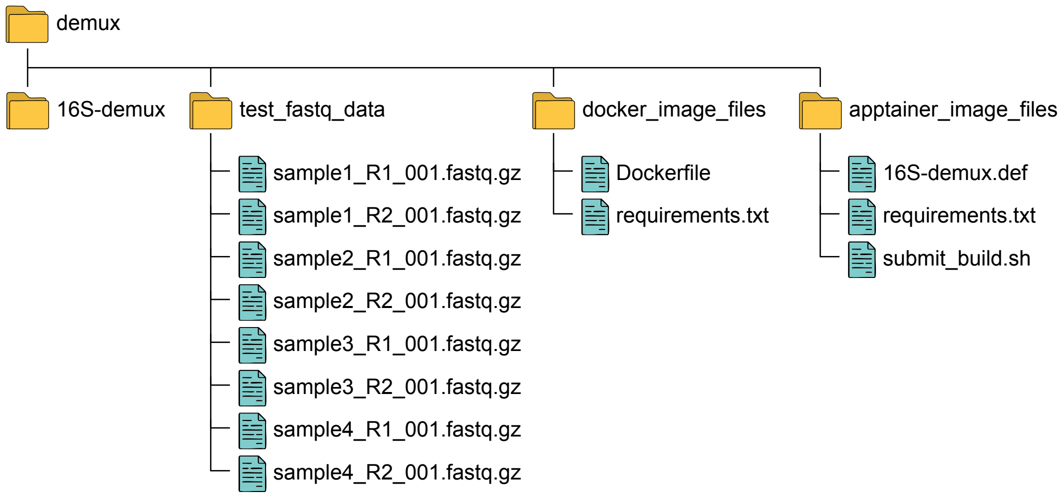


**Figure S1: Directory structure of the CUPID-seq demultiplexing package.**

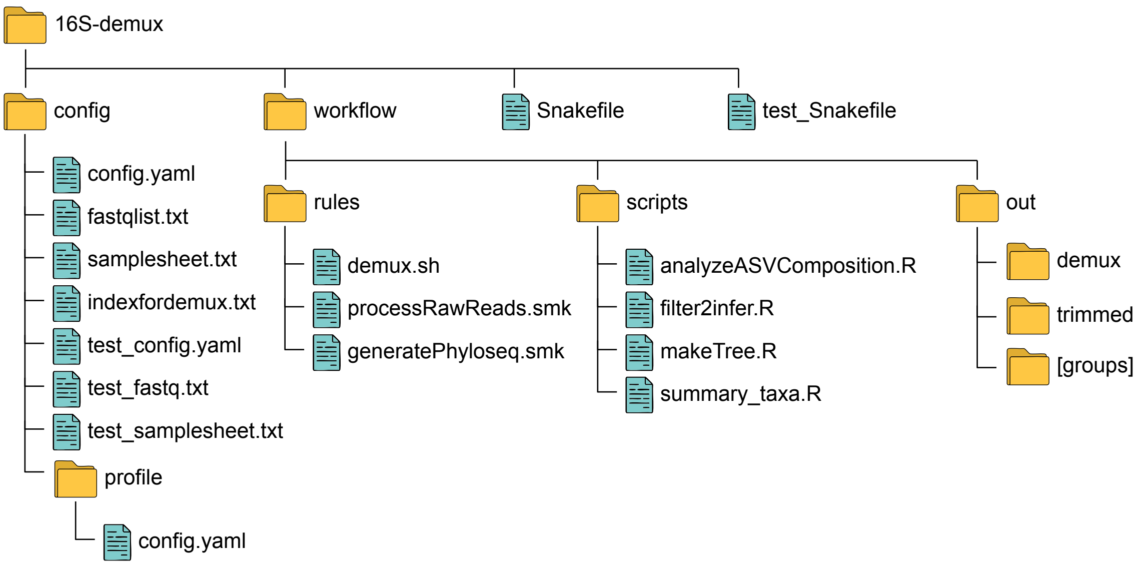


**Figure S2: Internal directory structure of the 16S-demux workflow.**

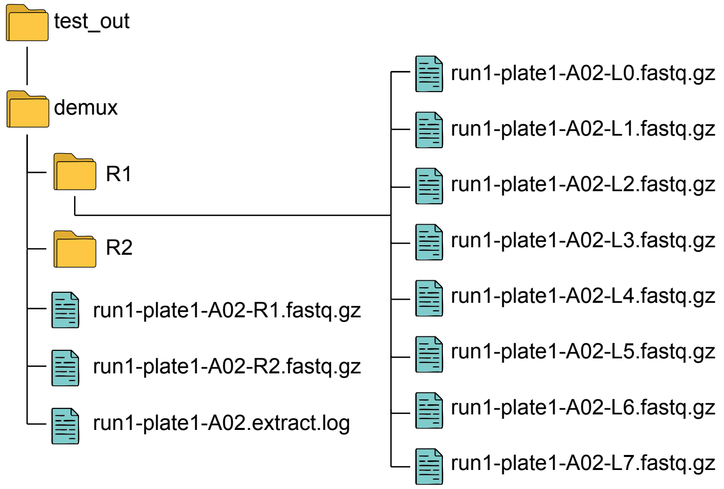


**Figure S3: Example directory structure and output files generated during Round–1 demultiplexing.**

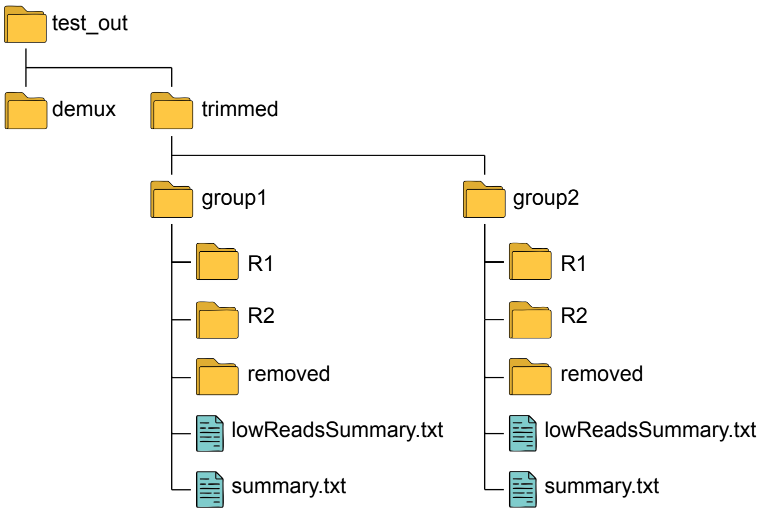


**Figure S4: Example directory structure and output files generated by the test workflow.**

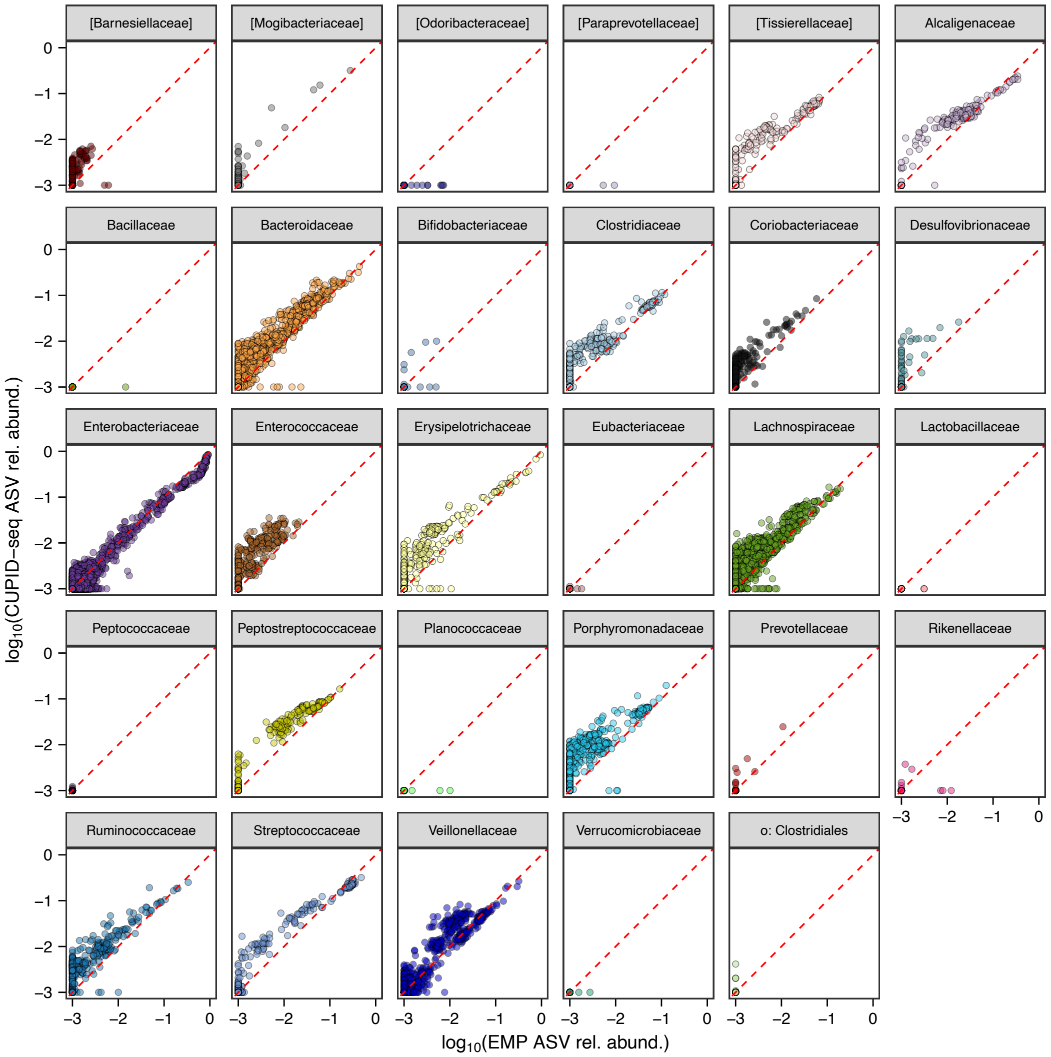


**Figure S5: Family-specific amplification differences between EMP and CUPID-seq protocols.** Correlation of $\log_{10}$-transformed ASV relative abundances from **Fig. 3a**, grouped by taxonomic family. Deviations from the diagonal indicate family-level differences in amplification efficiency between protocols.

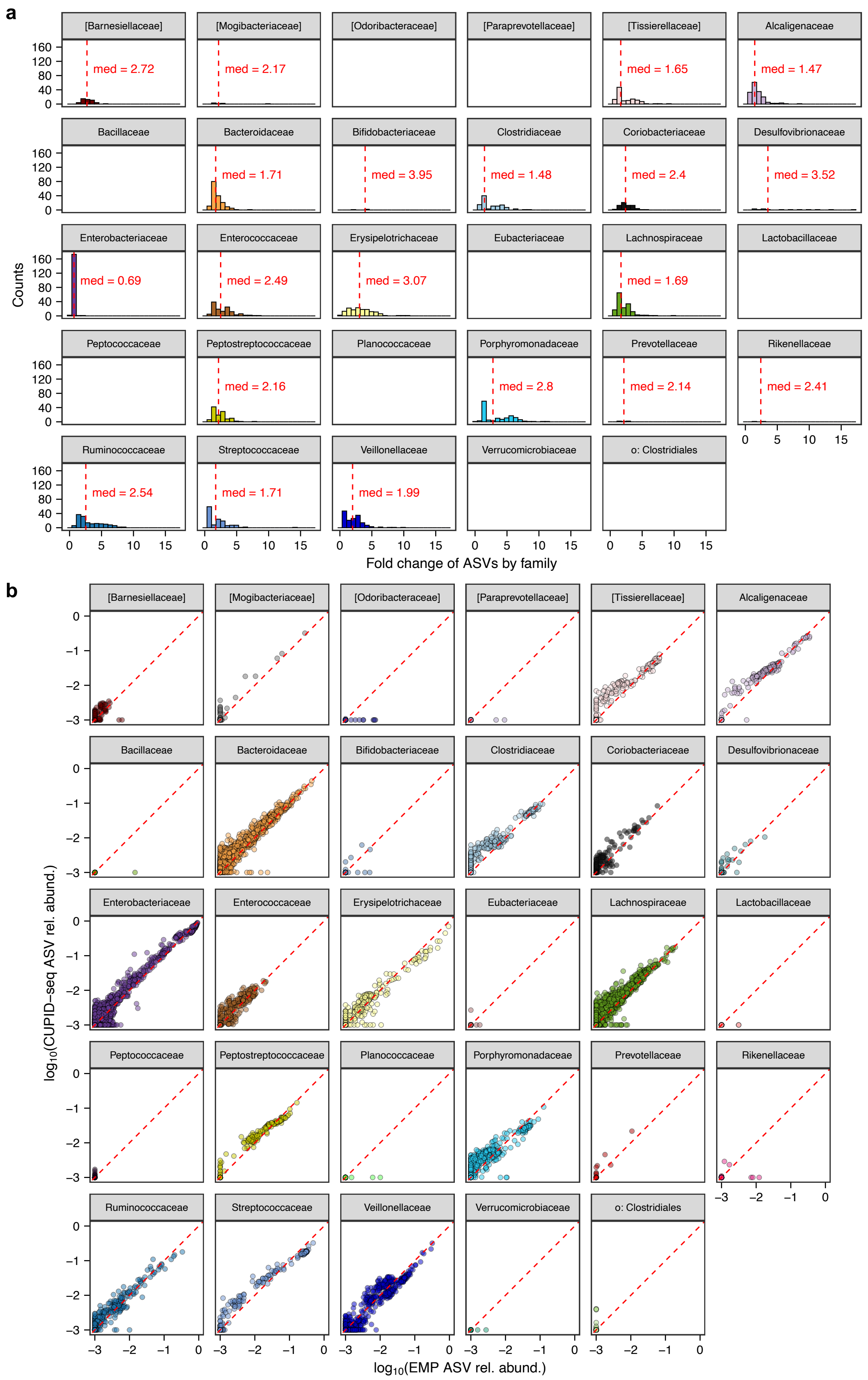
a
b
[Barnesiellaceae]
[Mogibacteriaceae]
[Odoribacteraceae]
[Paraprevotellaceae]
[Tissierellaceae]
Alcaligenaceae
Bacillaceae
Bacteroidaceae
Bifidobacteriaceae
Clostridiaceae
Coriobacteriaceae
Desulfovibrionaceae
Enterobacteriaceae
Enterococcaceae
Erysipelotrichaceae
Eubacteriaceae
Lachnospiraceae
Lactobacillaceae
Peptococcaceae
Peptostreptococcaceae
Planococcaceae
Porphyromonadaceae
Prevotellaceae
Rikenellaceae
Ruminococcaceae
Streptococcaceae
Veillonellaceae
Verrucomicrobiaceae
o: Clostridiales
med = 2.72
med = 2.17
med = 1.65
med = 1.47
med = 1.71
med = 3.95
med = 1.48
med = 2.4
med = 3.52
med = 0.69
med = 2.49
med = 3.07
med = 1.69
med = 2.16
med = 2.8
med = 2.14
med = 2.41
med = 2.54
med = 1.71
med = 1.99
Counts
Fold change of ASVs by family
$\log_{10}$(CUPID-seq ASV rel. abund.)
$\log_{10}$(EMP ASV rel. abund.)

**Figure S6: Correcting family-specific amplification differences improves agreement between EMP and CUPID-seq measurements.** (a) Distribution of fold changes in ASV relative abundance between the EMP and CUPID-seq measurements from the dataset shown in **Fig. 3a**, grouped by taxonomic family. Median fold changes for each family are indicated in red. (b) Correlation of $\log_{10}$-transformed relative abundances after rescaling ASVs according to family-level fold-change estimates derived from **Fig. 3a** and re-normalizing abundances within each sample. ASVs are shown separately for each taxonomic family.

### Methods

#### Genomic DNA (gDNA) extraction

gDNA was extracted from 50 µL of bacterial cultures using the DNeasy UltraClean 96 Microbial Kit (Qiagen, cat. no. 10196-4) according to the manufacturers' protocols.

#### 16S rRNA gene V4 sequencing using Earth Microbiome Project (EMP) primers

For amplification of the V4 region of the 16S rRNA gene, 1 µL of extracted gDNA was added to a 25 µL PCR containing EMP 515F/806R primer pairs[1] (0.4 µM final concentration) and AccuStart II PCR SuperMix (Quantabio, cat. no. 95137-100). PCR conditions were: 94 °C for 3 min; 35 cycles of 94 °C for 45 s, 50 °C for 60 s, and 72 °C for 90 s; followed by 72 °C for 10 min. Following amplification, 5 µL of each PCR product were pooled directly. The pooled library was cleaned and concentrated using the Macherey-Nagel NucleoSpin® Gel and PCR Clean-up Mini Kit (Fisher, cat. no. 740609) and sequenced using 300-bp paired-end reads on a MiSeq platform (Illumina).

#### Full-length 16S rRNA gene sequencing

Extracted gDNA was first cleaned using the OneStep PCR Inhibitor Removal Kit (Zymo Research, cat. no. D6030) and submitted to Zymo Research for library preparation and sequencing. Full-length 16S rRNA genes were amplified using primers 27F (5′-AGAGTTTGATCMTGGCTCAG-3′) and 1492R (5′-TACGGYTACCTTGTTAYGACTT-3′) Libraries were sequenced on a PacBio Sequel IIe system using SMRT Cell 8M chemistry (PacBio).

### References


1. Caporaso, J. G. *et al.* Moving pictures of the human microbiome. *Genome Biol.* **12**, R50 (2011).
